# Developmental oligodendrocytes regulate brain function through the mediation of synchronized spontaneous activity

**DOI:** 10.1101/2024.05.06.590880

**Authors:** Ryo Masumura, Kyosuke Goda, Mariko Sekiguchi, Naofumi Uesaka

**Affiliations:** Department of Cognitive Neurobiology, Graduate School of Medical and Dental Sciences, Institute of Science Tokyo, Tokyo 113-8510, Japan

## Abstract

Synchronized spontaneous neural activity is a fundamental feature of developing central nervous systems and is thought to be essential for proper brain development. However, the mechanisms that regulate this synchronization and its long-term impact on brain function remain unclear. Here, we identify a previously unrecognized role of oligodendrocytes in orchestrating synchronized spontaneous activity during a critical developmental window, with lasting consequences for adult behavior. Using oligodendrocyte-specific genetic manipulation in the mouse cerebellum, we demonstrate that oligodendrocyte deficiency during early postnatal development, but not after weaning, disrupts the synchronization of Purkinje cell activity both during development and in adulthood. The early disruption produced persistent deficits in cerebellar-dependent behaviors, including anxiety, sociality, and motor function. Optogenetic re-synchronization in adulthood restored motor and social functions but not anxiety-like behavior, demonstrating that reduced Purkinje cell synchrony specifically drives the motor and social impairments. Our findings establish a causal link between developmental oligodendrocyte-regulated neural synchrony and the emergence of complex brain functions, which depend on the proper developmental trajectory necessary for driving brain function.

## Introduction

Even before sensory inputs are fully established, immature animals can respond to environmental cues and generate diverse outputs, implying that complex neural networks are pre-organized. This organization relies on robust developmental processes, among which spontaneous activity plays a central role in shaping neural circuits. During early postnatal development, spontaneous activity is highly correlated across neural populations in various species and systems (Kirkby *et al*., 2013; Ackman & Crair, 2014). The widespread occurrence of correlated spontaneous activity suggests a universal principle of brain maturation, facilitating the formation of precise sensory maps and large-scale networks required for sensory processing and behavioral output.

In the developing central nervous system, correlated spontaneous activity initially manifests as synchronized patterns in localized or extensive regions (Adelsberger *et al*., 2005; Good *et al*., 2017; Babola *et al*., 2018; Mizuno *et al*., 2018; Murakami *et al*., 2022; Fujimoto *et al*., 2023). These patterns gradually transition to desynchronized states, as observed in the cortex, cerebellum, and olfactory bulb (Good *et al*., 2017; Mizuno *et al*., 2018; Fujimoto *et al*., 2023). Despite extensive documentation of these patterns, the mechanisms that generate synchronized activity and their functional significance remain poorly understood. Oligodendrocytes are strong candidates for regulating synchronized spontaneous activity because they control myelination and conduction velocity (Pajevic *et al*., 2014; Pajevic *et al*., 2023). Myelin formation continues throughout life, but the majority occurs during postnatal development in many brain regions (de Faria *et al*., 2021). However, how developmental oligodendrocytes contribute to brain maturation remains largely unknown.

Here we tried to determine the causal relationships among developmental oligodendrocytes, synchronized spontaneous activity, and brain function in the mouse cerebellum. The developing cerebellum is a suitable model, as the developmental trajectory of synchronized activity is well characterized (Good *et al*., 2017). To manipulate oligodendrocytes during development, we used an adeno-associated virus (AAV) vector with high oligodendrocyte specificity, enabling precise spatiotemporal reduction of these cells in the developing cerebellum. We found that reducing oligodendrocytes during the second to third postnatal week disrupted Purkinje cell (PC) synchrony. Behavioral analyses further revealed that perturbing developmental oligodendrocytes and synchronized activity led to lasting impairments in adult cerebellar function, including anxiety-like behaviors, reduced social interactions, and impaired motor coordination. Optogenetic re-synchronization in adulthood restored motor and social functions but not anxiety-like behavior. These results suggest that oligodendrocytes act as key regulators of synchronized spontaneous activity and are essential for the proper development of brain function.

## Materials and methods

### Experimental animals

This study was conducted under the recommendations of the Regulations for Animal Experiments and Related Activities at Tokyo Medical and Dental University. All animal experiment were approved by the Committees on Gene Recombination Experiments and Animal Experiments of Tokyo Medical and Dental University. All animals were group housed and maintained on a 12hr:12hr light dark cycle. All efforts were made to reduce the number of animals used and to minimize the suffering and pain of animals. WT mice (C57BL6N, 6 days -16 weeks, Sankyo labo service) were used in this study.

### Plasmids

The human MAG gene promoter sequences were amplified by PCR from human genomic DNA using primer pairs (Table. 1) and inserted into sites of hSyn promoter in AAV-U6-sgRNA-hSyn-mCherry (a gift from Alex Hewitt, Addgene plasmid # 87916) (AAV-hMAG-mcherry). hSyn promoter was removed by ApaI and BaMH1. The promoter was cloned using the Gibson Cloning Assembly Kit (New England BioLabs) following standard procedures. For AAV-hMAG-DTA, we amplified the DTA coding sequence from the pAAV-mCherry-flex-dtA (a gift from Naoshige Uchida, Addgene plasmid # 58536) using primer pairs (Table. 1) and cloned it into AAV-MAG-mcherry. The sequences of jGCaMP7f gene and jGCaMP7s were amplified by PCR from pGP-AAV-syn-jGCaMP7f-WPRE and pGP-AAV-syn-jGCaMP7s-WPRE (a gift from Douglas Kim & GENIE Project, Addgene plasmid #104488 and #104487) and inserted into sites of GFP in pAAV/L7-6-GFP-WPRE (a gift from Hirokazu Hirai, Addgene plasmid # 126462) (pAAV-L7-6-jGCaMP7f and pAAV-L7-6-jGCaMP7s) and into sites of GFP in pAAV-TRE-EGFP (a gift from Hyungbae Kwon, Addgene plasmid # 89875) (pAAV-TRE-jGCaMP7f and pAAV-TRe-jGCaMP7s). For AAV-TRE-Kir2.1-T2A-GFP, we amplified the sequences of Kir2.1-T2A-GFP from the pAAV hSyn FLEx-loxp Kir2.1-2A-GFP (a gift from Kevin Beier & Robert Malenka, Addgene plasmid # 161574), and cloned it into sites of GFP in pAAV-TRE-EGFP. For AAV-TRE-jGCaMP7f-T2A-Kir2.1, we amplified the sequences of jGCaMP7f and Kir2.1 from the pGP-AAV-syn-jGCaMP7f-WPRE and pAAV hSyn FLEx-loxp Kir2.1-2A-GFP, and cloned it into sites of GFP in pAAV-TRE-EGFP. For pAAV-TRE-rsChRmine-oScarlet, we amplified the sequences of rsChRmine-oScarlet from the pAAV -CaMKIIa-rsChRmine-oScarlet (a gift from Karl Deisseroth, Addgene plasmid # 183522) (Kishi *et al*., 2022), and cloned it into sites of GFP in pAAV-TRE-EGFP. For pAAV-L7-6-tTA, tTA was inserted into sites of GFP in pAAV/L7-6-GFP-WPRE.

### AAV packaging and injection

The AAVs were produced using standard production methods. HEK293 cells were transfected with Polyethylenimine. AAV9, PHPeB (Chan *et al*., 2017) and Olig001 (Powell *et al*., 2016) were used. Virus was collected after 120 h from both cell lysates and media. Purification Kit (Takara) was used for viral particle purification. All batches produced were in the range 10^10^ to 10^11^ viral genomes per milliliter. The procedure for viral vector injection has been modified from the previous lentivirus protocol(Uesaka *et al*., 2014). 1-1.5 µL of viral solution were injected into the cerebellum of C57BL/6N mice at 100 nl / min.

### Immunohistochemistry

Mice were perfused with 4% paraformaldehyde in 0.1 M phosphate buffer, and processed for parasagittal microslicer sections (100 μm in thickness). After permeabilization and blockade of nonspecific binding, the following antibodies were applied for 2 days at 4 °C: an antibody against Car8 (Car8-GP-Af500 or Car8-Go-Af780, diluted 1:300, Nittobo Medical) to visualize PCs; an antibody against S100b (S100b-GP-Af630, 1:300, Nittobo Medical); an antibody against Iba (Wako 019-19741, 1:1000), an antibody against RFP (390004, 1:1000, Synaptic Systems or PM005, 1:500, MBL), an antibody against MBP (NBP2-50035, 1:1000, Novus), an antibody against ASPA (ABN16986, 1:1000, Merck), an antibody against vGluT2 (MSFR106290, 1:300, Nittobo Medical), and/or an antibody against GFP (#06083-05, 1:1000, Nacalai Tesque). After incubation with secondary antibodies (an anti-rat Alexa Fluor 488, an anti-guinea pig Cy3, an anti-goat Alexa Fluor 488, an anti-goat Alexa Fluor 647, an anti-mouse Cy5, an anti-rabbit Cy3, an anti-rabbit Alexa Fluor 488), the immunolabeled sections were washed and then examined under a fluorescent microscope (BZ-X700 or BZ-X800, Keyence).

### In vivo calcium imaging

Animals were put on a warm blanket and anesthetized with a mixture of midazolam (4 mg/kg body weight (BW)), butorphanol (5 mg/kg BW), and medetomidine (0.3 mg/kg BW). The depth of anesthesia was constantly monitored by observing pinch-induced forelimb withdrawal reflex. The skin and muscles over the skull were removed and a metal plate was fixed with dental acrylic cement over the lobule 5-7 of the cerebellar vermis. A craniotomy of 3 mm diameter was made and the dura mater was carefully removed. A sterile circular 3mm diameter glass coverslip (#CS01078 3mm diameter, Matsunami) was put directly on the dura mater and was secured in place with surgical adhesive (Alon-Alpha A, Sankyo). A custom-made metal headplate with a 5 mm circular imaging well was fixed to the skull over the cranial window with dental cement (Super-Bond, Sun-medical, Japan). The head of the mouse was immobilized by attaching the head plate to a custom-made stage. We then administered atipamezole (0.3mg/kg, anti-sedan) (100 μg/g of body weight, intraperitoneal injection) for anesthesia reversal. After a 10 - 15 minutes waiting period, in vivo calcium imaging was performed using an Olympus MVX10 fluorescence microscope or a two-photon microscope (Nikon, AX R MP) equipped with a ×16 objective lens (Nikon, CFI75 LWD 16X W) and an ultrafast laser (Axon 920-2 TPC). During the recordings, the animals were laid on a warm blanket to keep the body temperature at 37°C. Laser power for the two-photon microscope was maintained <20 mW at the sample. Images were obtained in a 2D plane (2048 x 2048 pixels) using Andor SOLIS software (Oxford Instruments, frame rate 5 Hz) or in a 2D plane (512 × 512 pixels) using Nikon NIS-Elements software (frame rate, 1 Hz) for the two photon microscope. Data were saved as TIFF files.

We resized raw calcium imaging videos (TIFF format) to a resolution of 512 x 512 pixels with an 8-bit depth using ImageJ (JAVA-based processing program, USA). We cropped out the unwanted areas. Regions of interest (ROIs) corresponding to the active PC dendrites were detected using the Suite2p software (Marius Pachitariu, GitHub). This software executed a two-dimensional phase-correlation-based non-rigid image registration in the XY plane, extracted the ROIs considered as PC dendrites, and calculated the fluorescence intensity changes in each ROI. ROIs were extracted using the parameters shown in Tanigawa et al., 2024 (Tanigawa *et al*., 2024). We exported these data as MAT files. The MAT files were imported into MATLAB R2023b (MathWorks, USA) for further analysis using custom-made MATLAB routines. The fluorescence intensity (F) was converted to ΔF/F0 wave, expressed as ΔF/F0 = (F - F0)/(F0), where F0 is the baseline fluorescence in the absence of calcium transients. F0 was defined as follows. First, we calculated a threshold as mean + 1 SD of the F wave of each ROI, and then the F wave below the threshold was averaged to obtain F0. Parameters such as amplitude, frequency, half-width, and area of each calcium transient were quantified by the custom-made MATLAB routines. The correlation coefficient among all ROI pairs was calculated as previously described (Tsutsumi *et al*., 2015). The resulting correlation matrix captured relationships among all ROI pairs in the dataset.

### Behaviors

Male mice from 2 to 3 months of age 6-7 days after AAV injection were used for behavioral tests. Before the experiments, the mice were habituated in the testing areas for at least 30 min. Mice behavior was recorded through the video camera set in the experimental room. The open field test was performed in a 50 cm x 50 cm x 40 cm (W x D x H) open field box for 10 min. Total distance mice traveled and the time in the center and peripheral regions were automatically analyzed by the ImageJ software (MouBeAT) (Bello-Arroyo *et al*., 2018).

The sociality test was performed in the open field apparatus (50 x 50 x 40 cm, W x D x H). The test consisted of a 10-min habituation session and a 10-min test session. In the habituation session, mice were allowed to explore the open field freely. In the test session, two quadrangular cages with slits (8 cm x 8 cm, 18 cm high) were placed in two adjacent corners. One cage contained a novel mouse (8-week-old male C57BL6N) and the other was a mouse doll, and a test mouse was first placed in the outer regions of open field arena at the longest distance from these cages. The times spent around the two cages areas (12 cm in diameter) were analyzed automatically with the ImageJ software (MouBeAT). Social preference was assessed by the following equation: Sociality index = (T_mouse_)/(T_doll_) where T_mouse_ and T_doll_ are the staying time around the mouse cage and that around the doll cage, respectively.

To assess motor coordination and motor learning, mice were subjected to a rotarod test. Mice were placed on a resting state rotarod (model LE8205, Panlab) for three consecutive days with two trials per day spaced 10 min apart (30 min and 40 min after CNO injection). For each trial, the rotarod was accelerated linearly from 4 rpm to 40 rpm over 300 s. Latency to fall from the start of rotation was measured.

### Optogenetic stimulation

To manipulate PC synchrony, we expressed a red-shifted channelrhodopsin variant selectively in PCs of control and DTA mice by stereotaxic injection of pAAV-TRE-rsChRmine-oScarlet and pAAV-L7-6-tTA into the cerebellar vermis 2 weeks before behavior tests. Mice were subjected to behavioral assays at P60-70. For light delivery, we used a wireless optogenetic system (Teleopto; Bio Research Center Co., Ltd., Aichi, Japan) equipped with a brain-surface LED probe (TeleLP-c, 590 nm). After making a small midline skin incision, the LED probe was placed directly above the cerebellar vermis on the intact skull and secured with dental cement (Super-Bond C&B). The cement was coated with black nail polish to minimize light leakage. This approach allowed stable illumination of the cerebellar surface without penetrating the tissue. Probe placement was verified after the completion of behavioral experiments. During behavioral testing, 590-nm light pulses (10 ms, 1 Hz) were delivered throughout the task sessions using a Teleopto Remote Controller via an infrared emitter. Stimulation was applied during open-field, three-chamber social interaction, and rotarod assays. Illumination parameters were selected to evoke population-wide synchronous PC spiking while minimizing nonspecific effects on locomotor activity.

### Statistical analysis

All data are presented as mean ± SEM. Statistical significance was assessed by Mann-Whitney U test for the comparison of two independent samples. To compare the two different categorical independent samples on one dependent variable, Two-way ANOVA with post-hoc test (Bonferroni correction) was used as indicated in the text. For datasets in which multiple measurements were obtained from the same animal (e.g., multiple fields of view or recordings), statistical dependence within animals was accounted for by performing nested ANOVA or nested t-tests, treating animal identity as the nesting factor. This ensured that repeated measurements within the same animal were not treated as independent replicates. Because the nested models yielded comparable effect sizes to the Mann–Whitney tests, we have retained the mean ± SEM for ease of comparison with prior literature but now also report all values for each mouse in Table 1. Statistical analysis was conducted with GraphPad Prism program. Differences between groups were judged to be significant when p values were smaller than 0.05. *, **, ***, **** and ns represents p < 0.05, p < 0.01, p < 0.001, p < 0.0001 and not significant, respectively.

## Results

### Developmental oligodendrocytes regulate synchronized spontaneous activity

We tested the role of developmental oligodendrocytes in regulating synchronized spontaneous activity and brain function. We generated an oligodendrocyte-specific AAV (OL-AAV) containing a human myelin associated glycoprotein (hMAG) promoter and an oligodendrocyte-tropic capsid (Figure 1A). We focused on the developing mouse cerebellum in which synchronized spontaneous activity and its transition were observed during postnatal development (Good *et al*., 2017). Myelination in the cerebellum appeared by postnatal day 5 (P5) and P7 (Figure supplement 1A-B).

**Figure 1.**
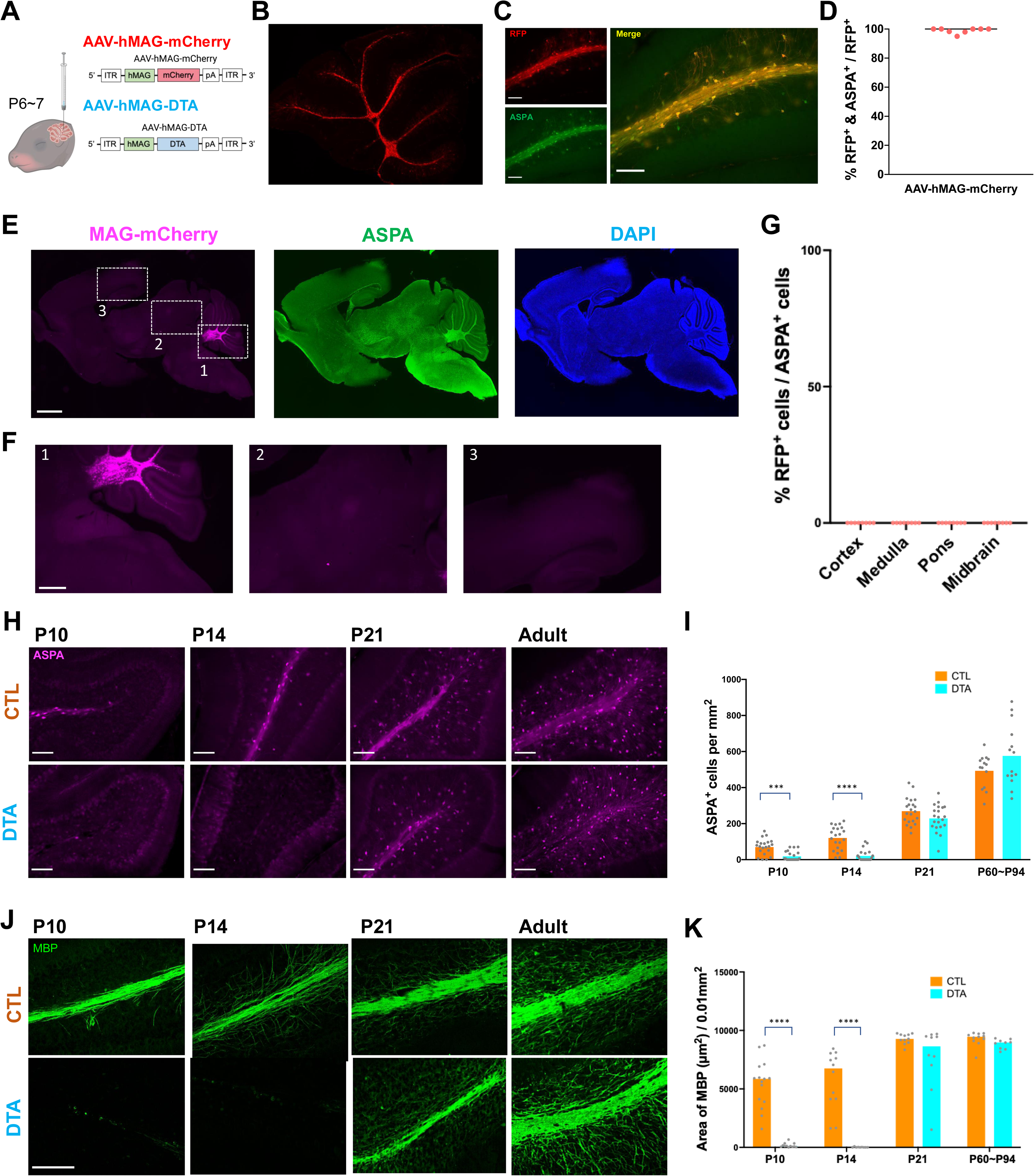
DTA expression by oligodendrocyte-specific AAVs deletes oligodendrocytes in the mouse cerebellum during the specific developmental period. **A**, Diagram illustrating the injection of AAV-hMAG-mCherry and AAV-hMAG-DTA into the mouse cerebellum at early postnatal days (P6-7). **B**, Fluorescence microscopy image showing mCherry expression (red) in the section of cerebellum at P14 following AAV-hMAG-mCherry injection at P7. Scale bar, 300 µm. **C**, Specificity of mCherry expression (red) in ASPA-positive oligodendrocytes (green) at P14 by AAV-hMAG-mCherry injection at P7. Scale bar, 100 µm. **D**, Scatter plot graph depicting the percentage of mCherry and ASPA double-positive cells among mCherry-positive populations (8 regions from 2 mice). **E**, Low-magnification image showing the distribution of MAG-mCherry expression across sagittal brain sections at P14 after AAV-hMAG-mCherry injection at P7. Expression is confined to the cerebellum and absent in extracerebellar regions. Scale bar, 500 µm. **F**, Higher-magnification views of regions indicated in panel E. Robust mCherry expression is observed in the cerebellum (1), whereas no detectable expression is found in the midbrain (2) or cerebral cortex (3). Scale bar, 200 µm. **G**, Quantification of the regional specificity of MAG-mCherry expression. Scatter plots indicate the percentage of mCherry-positive cells detected in the cortex, medulla, pons, and midbrain (4 regions from 2 mice). There was no mCherry-positive cell in these regions. **H**, Sequential visualization of oligodendrocyte reduction over time, indicated by ASPA staining in cerebellar sections from control (AAV-hMAG-mCherry) and DTA-treated (AAV-hMAG-DTA) mice at P10, P14, P21, and P78. Scale bar, 100 µm. **I**, Quantification of ASPA-positive cell density at each stage (from 4 mice per group per each stage). Bars and dots indicate mean and data from individual fields of view, respectively. Adulthood data were collected at P60-P94. **** p < 0.0001 (Nested t-test). **J,** Representative images of MBP immunostaining (green) in the cerebellar white matter of control and DTA-treated mice at P10, P14, P21, and adulthood (P60). Scale bar, 100 µm. **K**, Quantification of MBP-positive bundle area at each stage (from 4 mice per group per stage). Bars and dots indicate mean and data from individual fields of view, respectively. p < 0.05, **** p < 0.0001 (Nested t-test).

Injection of OL-AAVs expressing mCherry into the cerebellum of P7 mice resulted in robust and selective labeling of cerebellar oligodendrocytes by P21, with >98% of mCherry-positive cells co-expressing the mature oligodendrocyte marker ASPA (Figure 1B-D). In contrast, mCherry was not detected in Olig2-positive but ASPA-negative immature oligodendrocyte-lineage cells, confirming specific targeting of mature oligodendrocytes (Figure supplement 1C-D). mCherry expression was restricted to the cerebellum and was not detected in extracerebellar regions (Figure 1E-G), confirming the regional specificity of this approach.

To selectively ablate oligodendrocytes, we injected OL-AAVs carrying Diphtheria toxin A (DTA) into the cerebellum of P7 mice. This manipulation resulted in a significant reduction of oligodendrocyte density (ASPA^+^ cells) and myelin coverage (MBP staining) by P10 and P14 (Figure 1H-K and Figure supplement 1E). The density and coverage returned to control levels by P21 and remained normal in adulthood, indicating a transient but robust disruption of oligodendrocytes and myelination during a defined developmental window. This temporal profile provided the opportunity to test how developmental oligodendrocytes regulate PC activity synchrony and its maturation. Despite the transient loss of mature oligodendrocytes, gross cerebellar morphology remained indistinguishable from controls (Figure supplement 1F-G).

To determine whether other cell types were affected (Mathis *et al*., 2003), we examined microglia, astrocytes, PCs, and molecular layer interneurons in DTA-ablated mice. IBA1 staining showed no differences in microglial density and IBA1 intensity between DTA and control mice at both of P14 and P21 (Figure supplement Fig. 2A-D). S100β staining showed no differences in Bergmann glia density and S100β intensity (Figure supplement Fig. 2E-H). Parvalbumin (PV) immunostaining demonstrated that PC density and dendritic arborization were preserved in DTA mice (Figure supplement Fig. 2I-K). PV-positive interneurons in the molecular layer were present at normal density in DTA mice (Figure supplement Fig. 2L). The same results were obtained at P21 (Figure supplement Fig. 2M-P). Together, these results indicate that transient developmental ablation of oligodendrocytes does not induce compensatory gliosis or neuronal loss.

We next tested whether oligodendrocyte deficit affected synchronized spontaneous activity in the cerebellar PC population (Figure 2A-B). Here, spontaneous activity refers to neural events occurring in the absence of external sensory stimulation. Developmental PCs display highly synchronized CF–driven activity in newborn mice and undergo progressive desynchronization as development advances (Good *et al*., 2017). In our imaging paradigm, regions of interest were restricted to the dendritic arbor of PCs where calcium signals are dominated by climbing-fiber (CF) synaptic inputs (Ramirez & Stell, 2016; Good *et al*., 2017). In vivo calcium imaging of PCs revealed a marked reduction in synchrony of spontaneous activity among PCs at P13-15 following oligodendrocyte reduction in DTA mice (Figure 2C-F). This effect was observed across both proximal and distal PC populations (Figure 2E-F), indicating a widespread disruption of neural activity synchronization. Notably, other parameters such as frequency, amplitude, and area of calcium transients remained unchanged (Figure 2G-I), suggesting a specific effect on PC activity synchrony rather than general neural activity.

**Figure 2.**
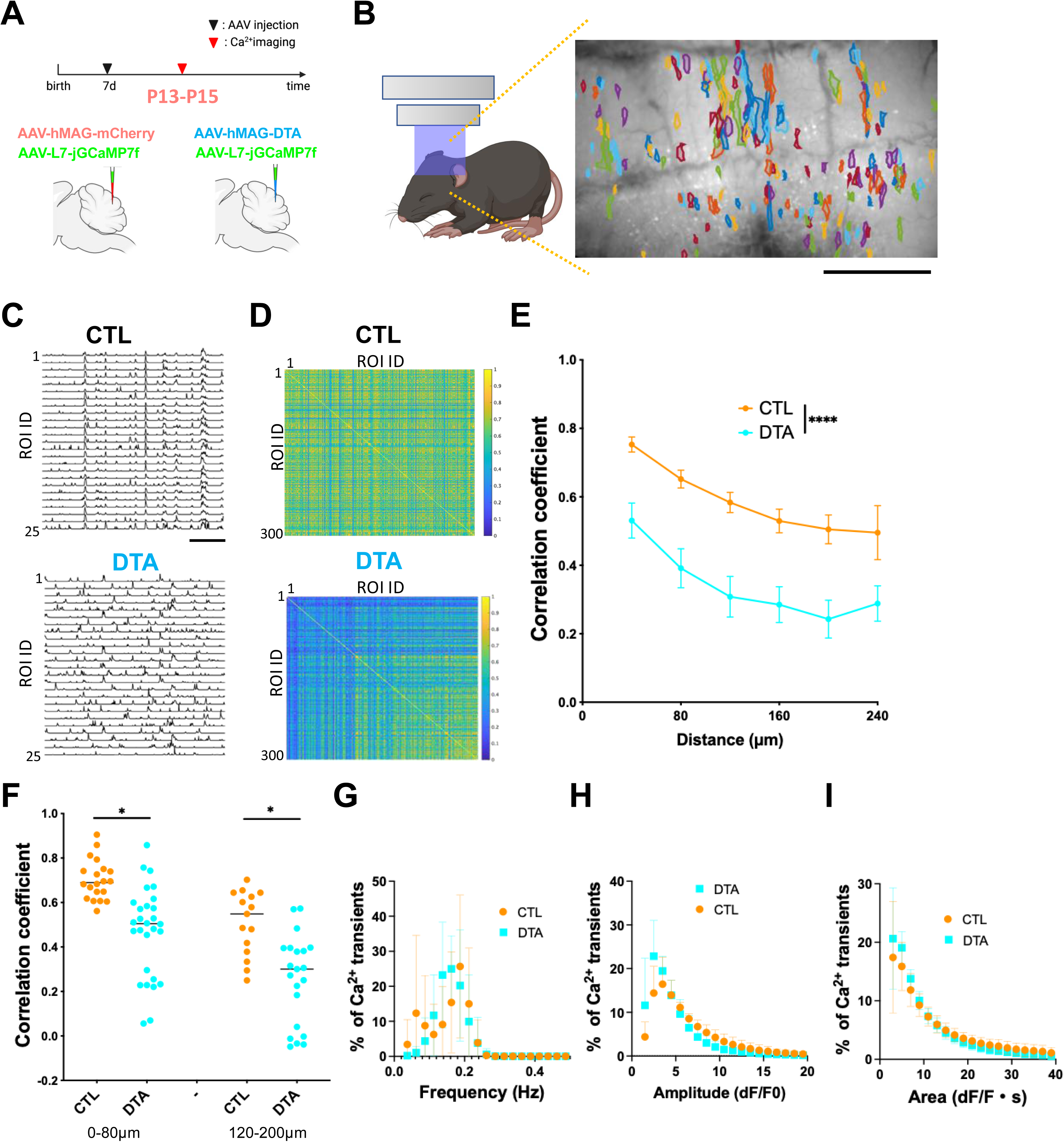
Cerebellar oligodendrocyte ablation reduces the synchrony of PC population activity during mouse cerebellar development. **A,** Experimental timeline showing AAV injection at P7 and in vivo calcium imaging at P13–P15 in control and DTA-expressing mice with jGCaMP7f expressed in PCs. **B**, Schematic of in vivo one photon calcium imaging and a representative frame from a field scan in lobule VII/VIII of a control mouse at P14 with analyzed ROIs. Scale bar, 0.5 mm. **C-F**, Decrease in correlation coefficients (CCs) indicating reduced synchrony of spontaneous activities among PC population at P13-15 following DTA-mediated oligodendrocyte ablation. **C**, Representative traces of spontaneous calcium transients (ΔF/F0) extracted from 25 regions of interest (ROIs) in the cerebellum of control and DTA mice at P14. Scale bar, 20 s. **D**, Correlation coefficient matrices of spontaneous calcium activity across 300 ROIs in control (top) and DTA (bottom) mice at P14. Color scale indicates the strength of pairwise correlations. **E**, Mean correlation coefficients between calcium traces from pairs of ROIs plotted against the distance of separation between pairs of ROIs in control (n = 5 mice) and AAV-hMAG-DTA-injected (n = 7 mice) mice at P13-15. Error bars indicate SEM. **** p < 0.0001 (two-way ANOVA). **F**, Scatter plot graph of CCs for ROI pairs, categorized by separation distances of 0–80 mm and 120–200 mm (analyzed from four mice per group). Lines and plots indicate mean and data from individual separation distances, respectively. Significant group differences were observed (* p < 0.05, Nested t-test, 4 mice for each). **G-I**, Frequency distributions of Frequency (G), Amplitude (H), and Area (I) of calcium transients in CTL (AAV-hMAG-mCherry) and DTA-treated (AAV-hMAG-DTA) mice at P13–15. Circles and squares indicate data from individual ROIs, and bars represent SEM. There was no significant difference between groups (two-way ANOVA).

Having found that developmental oligodendrocyte loss decreases PC synchrony, we next examined whether this effect originated from altered CF activity. We imaged CF population activity by expressing jGCaMP7f in inferior olive neurons (Figure 3A). Correlation analysis of CFs revealed that CF–CF synchrony, but not amplitude, frequency, area, width, was significantly reduced in DTA mice at P13-15 (Figure 3B–D), indicating that impaired CF coordination contributes to the reduction of PC synchrony. Electrophysiological analyses showed that parallel-fiber EPSCs and inhibitory synaptic inputs onto PCs were not significantly altered (Figure 3E–H), making it unlikely that reduced synchrony results from altered excitatory or inhibitory drive other than CF inputs. Together, these findings demonstrate that developmental oligodendrocytes are required to maintain CF-driven PC synchronization during a critical window of cerebellar maturation.

**Figure 3.**
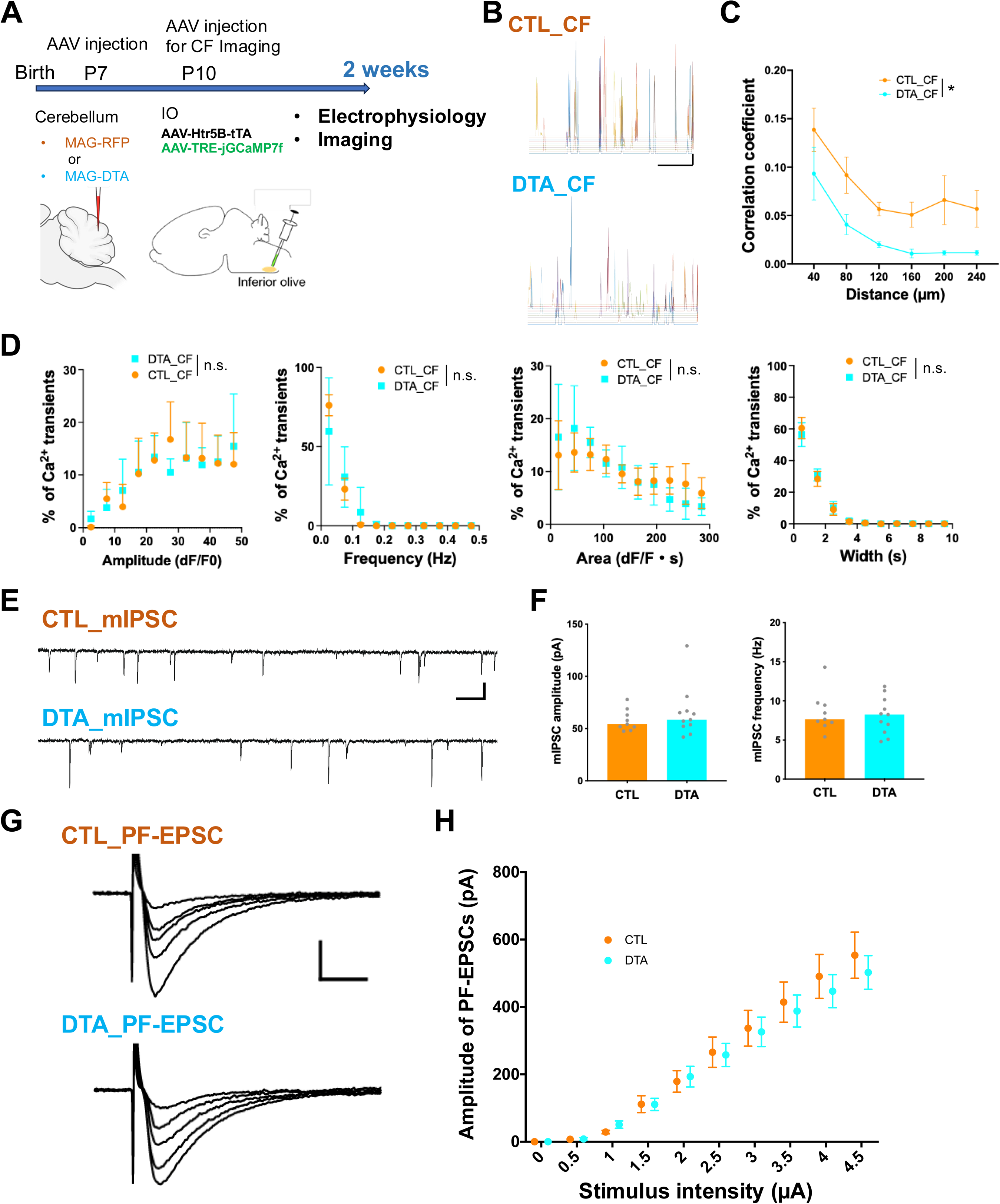
Developmental oligodendrocyte ablation reduces CF synchrony but does not alter PF excitatory or inhibitory synaptic inputs to PCs. **A**, Experimental scheme. AAV-hMAG-RFP or AAV-hMAG-DTA was injected into the cerebellum at P7, and AAV-TRE-jGCaMP7f was injected into the inferior olive at P10 to label CFs. CF activity was imaged in the cerebellar cortex at P13-15. **B**, Representative calcium traces of CFs from control (CTL) and DTA-treated mice at P14. Scale bars, 60 s, 5 dF/F0. **C**, Cross-correlation analysis showing reduced CF–CF synchrony in DTA mice compared to controls at P13-15. Mean correlation coefficients between calcium traces from pairs of ROIs plotted against the distance of separation between pairs of ROIs in control (n = 6 mice) and AAV-hMAG-DTA-injected (n = 5 mice) mice. Error bars indicate SEM. * p < 0.05 (two-). **D**, Frequency distributions of CF calcium transient properties, including amplitude, event frequency, Area, and width in CTL and DTA mice. No significant group differences were observed (two-way ANOVA). **E**, Representative traces of miniature inhibitory postsynaptic currents (mIPSCs) recorded from PCs in CTL and DTA mice at P14. Scale bars, 100 ms, 50 pA. **F**, Quantification of mIPSC amplitude and frequency showing no significant differences between groups (9 mice for control, 11 mice for DTA, Mann–Whitney U test). Bars indicate median, dots represent the data for individual animals. **G**, Representative traces of parallel fiber–evoked EPSCs (PF-EPSCs) recorded from PCs in CTL and DTA mice at P14. Scale bars, 5 ms, 200 pA. H, Input–output curves of PF-EPSCs demonstrating no significant differences in excitatory drive between groups at P13-15 (two-way ANOVA). Data are shown as mean ± SEM (5 mice for each).

### Developmental oligodendrocyte affects brain function

We next examined whether developmental oligodendrocytes and synchronized spontaneous activity affect adult brain function. We first investigated the long-term effects of oligodendrocyte reduction during cerebellar development on the synchrony of PC activity in the adult cerebellum. We selectively ablated oligodendrocytes in early postnatal mice and monitored PC activity as the animals matured. Developmental reduction of oligodendrocytes caused persistent hyposynchrony of PC activity in adulthood, whereas other parameters of PC activity remain unaffected (Figure 4). These findings suggest that disturbances in oligodendrocyte function during development have lasting consequences for brain function, underscoring the role of developmental oligodendrocytes in sustaining neuronal synchrony across the lifespan.

**Figure 4.**
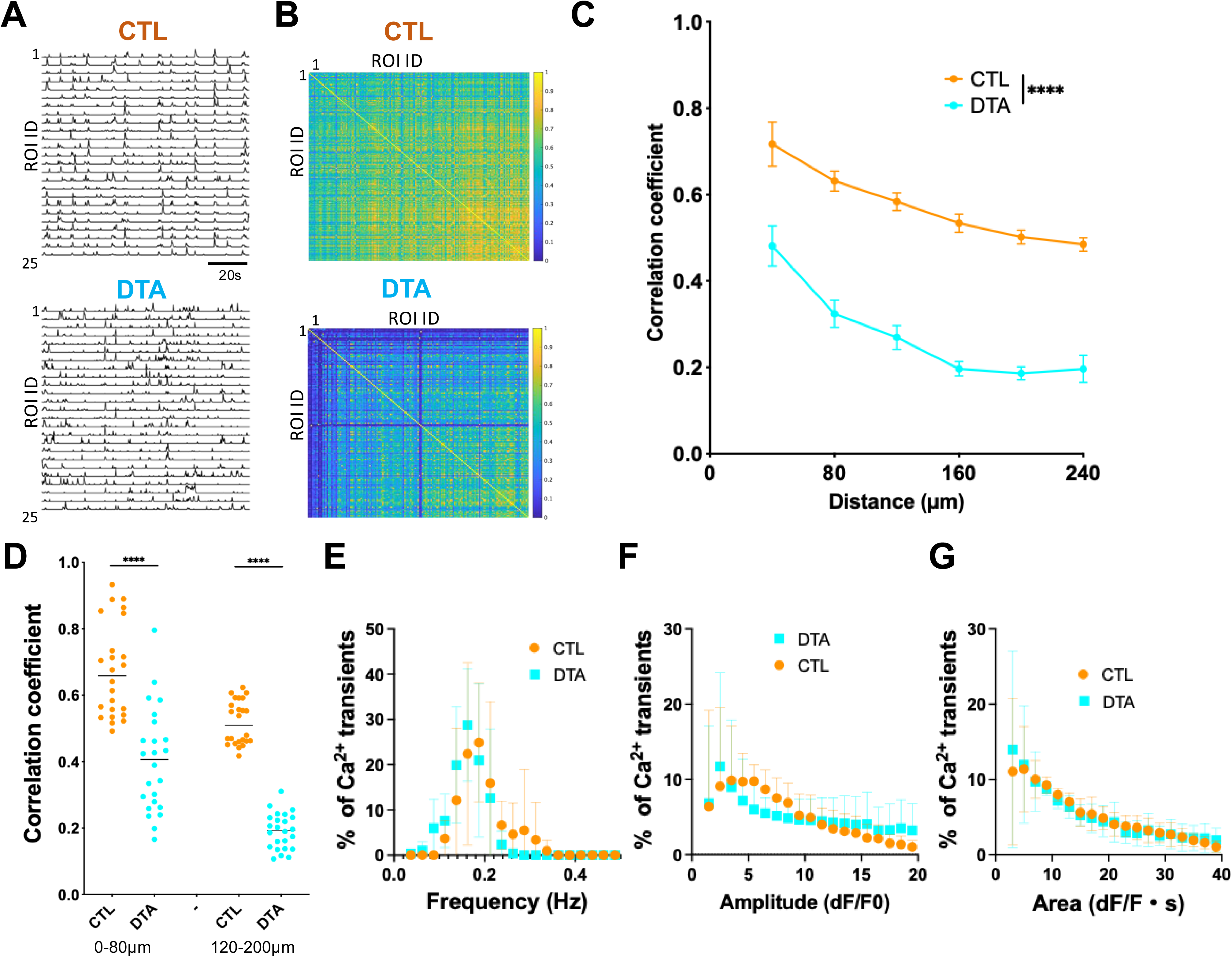
Oligodendrocyte ablation during postnatal development diminishes neuronal synchrony at adult stage. **A,** Representative traces of spontaneous calcium transients (ΔF/F0) extracted from 25 regions of interest (ROIs) in the cerebellum of control and DTA mice at P62. Scale bar, 20 s. **B**, Correlation coefficient matrices of spontaneous calcium activity across multiple ROIs in control (top) and DTA (bottom) mice at P62. Color scale indicates the strength of pairwise correlations. **C**, Mean correlation coefficients between calcium traces from pairs of ROIs plotted against the distance of separation between pairs of ROIs in control (n = 6 mice) and AAV-hMAG-DTA-injected (n = 6 mice) mice at P62-80. Error bars indicate SEM. **** p < 0.0001 (two-way ANOVA). **D**, Scatter plot graph of CCs for ROI pairs, categorized by separation distances of 0–80 mm and 120–200 mm (analyzed from four mice per group). Lines and plots indicate median and data from individual separation distances, respectively. Significant group differences were observed (**** p < 0.0001, Nested t-test, 4 mice for each). **E-G**, Frequency distributions of Frequency (E), Amplitude (F), and Area (G) of calcium transients in CTL (AAV-hMAG-mCherry) and DTA-treated (AAV-hMAG-DTA) mice at P62-80. Circles and squares indicate data from individual ROIs, and bars represent SEM. There was no significant difference between groups (two-way ANOVA).

Because developmental oligodendrocyte loss disrupted PC synchrony in adulthood, we next asked how this perturbation impacts brain function at the behavioral level. The cerebellum contributes not only to sensory-motor functions, including chewing, postural control, tone, movement, the coordination of balance, eye movements, and reflexes (Stoodley & Schmahmann, 2018; Algin *et al*., 2023), but also to cognitive, social, and emotional processes (Depping *et al*., 2018; Schmahmann, 2019; Van Overwalle *et al*., 2020). To examine roles of developmental oligodendrocytes in these brain function, we injected OL-AAVs with hMAG-mCherry or hMAG-DTA at P6-7 into distinct cerebellar compartments (left, center, right) (Figure supplement Fig. 3) corresponding to functional domains (Manni & Petrosini, 2004; Guell *et al*., 2018). Behavioral assessments in adulthood unveiled that oligodendrocyte reduction during development induced anxiety-like behavior specifically when targeted to the central cerebellar region, without altering distance travelled, underscoring the compartment-specific contribution to anxiety regulation (Figure 5A-C). Across all regions, oligodendrocyte reduction led to reduced social interaction and impaired motor coordination, as shown by reduced social approach to novel mouse and shorter latency to fall in the Rotarod test, respectively (Figure 5D-G). These data indicate that developmental oligodendrocytes are essential for mental states, sociality, and motor function in adult stages.

**Figure 5.**
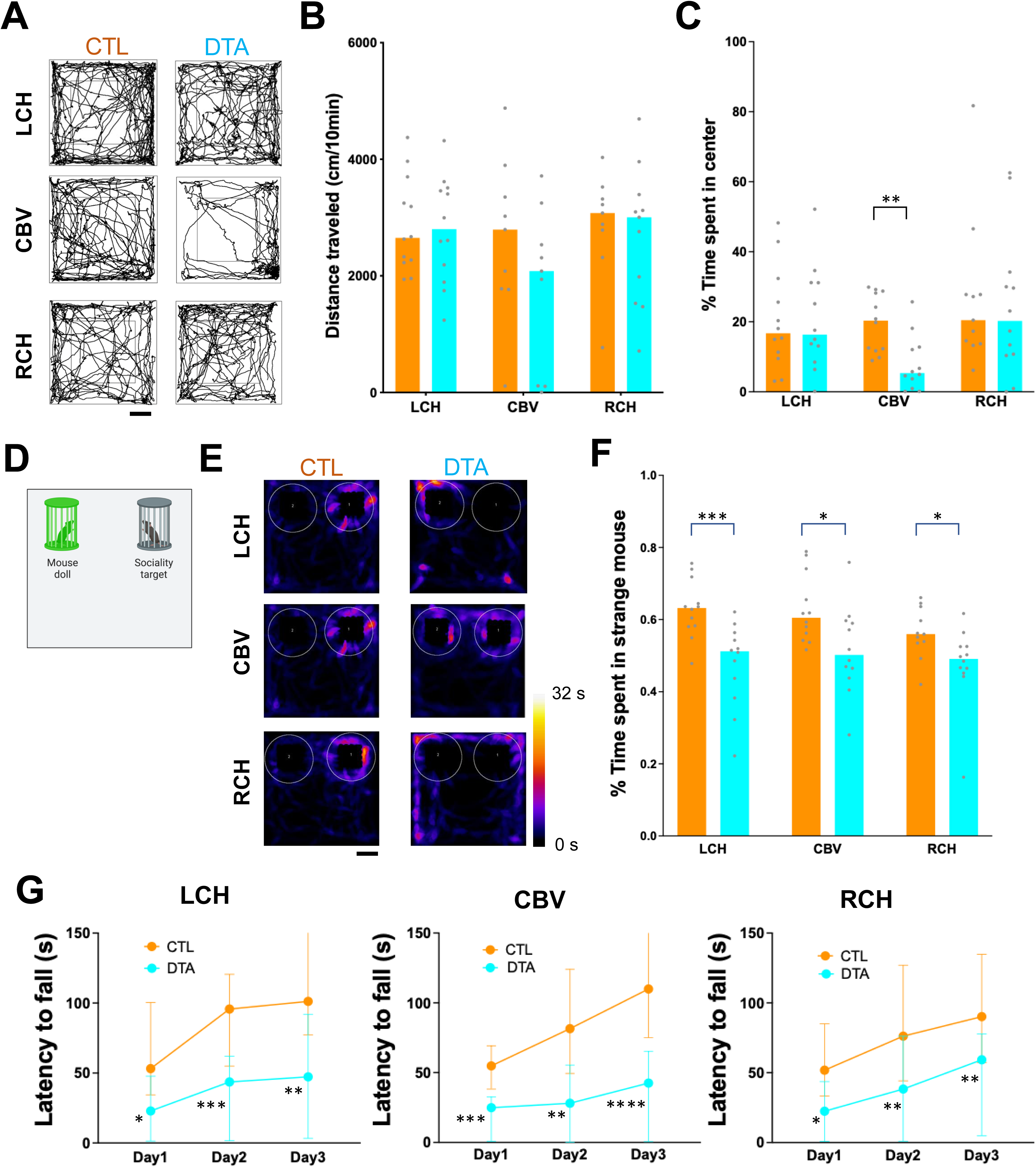
Oligodendrocyte reduction during development leads to behavioral abnormalities. **A-C,** Assessment of exploratory behavior and locomotor activity in control at P60 (CTL, orange, 12 mice for each injection site) and DTA-injected at P60 (DTA, cyan, 12 mice for each injection site) mice in a novel environment, with trajectories of mouse exploratory behavior (**A**), distance traveled (**B**), and time spent in center (**C**) for mice with the injection of AAVs into distinct compartments (left, center, right) of the cerebellum (LCH, CBV, RCH) at P60-80. Bars and dots indicate median and data from individual mice, respectively. **D-F,** Representative images and quantitative analysis of sociality in CTL and DTA mice with the injection of AAVs into distinct compartments (LCH, CBV, RCH). Diagram in (**D**) shows the method of sociality test; (**E**) displays the heatmap of mouse exploratory behavior, and (**F**) shows the percentage of time spent around strange mouse with data points representing individual mice at P60-80. Bars and dots indicate median and data from individual mice, respectively (CTL, 12 mice for each site, DTA, 12 mice for each site). **G,** Line graph showing the median of latency to fall in the rotarod test over a three-day period for CTL and DTA mice at P60-80, with separate graphs for mice with the injection of AAVs into distinct compartments (LCH, CBV, RCH). Error bars indicate 95 % CI. Statistical significance is indicated by asterisks (*p < 0.05, **p < 0.01, *** p < 0.001, **** p < 0.0001, Mann-Whitney U tests).

### Developmental oligodendrocytes during specific time windows affect brain functions

We next examined whether oligodendrocytes influence cerebellar function within specific developmental time windows. To address this, we administrated hMAG-DTA to the central cerebellum at postnatal week 3 (3W_DTA mice), shortly after weaning. In these mice, oligodendrocytes in the cerebellum were significantly reduced at 4 weeks (1 week after AAV injection) compared with control mice with only hMAG-mCherry, but returned to near-control levels by 2 months old (Figure 6A-C), indicating that the ablation was temporary. Calcium imaging of 3W_DTA mice at 4weeks old revealed no significant alteration for correlation coefficient and activity (Figure 6D-J). These results suggest that oligodendrocytes before and after the weaning exert distinct influences on PC activity. Behavioral assessments in adulthood of these mice uncovered no significant changes in anxiety or sociality (Figure 6K-O). However, 3W_DTA mice showed a subtle motor impairment as evidenced by shorter latency to fall at day3 in the Rotarod test (Figure 6P). These data suggest that oligodendrocytes before and after weaning make temporally distinct contributions to neural activity and brain function.

**Figure 6.**
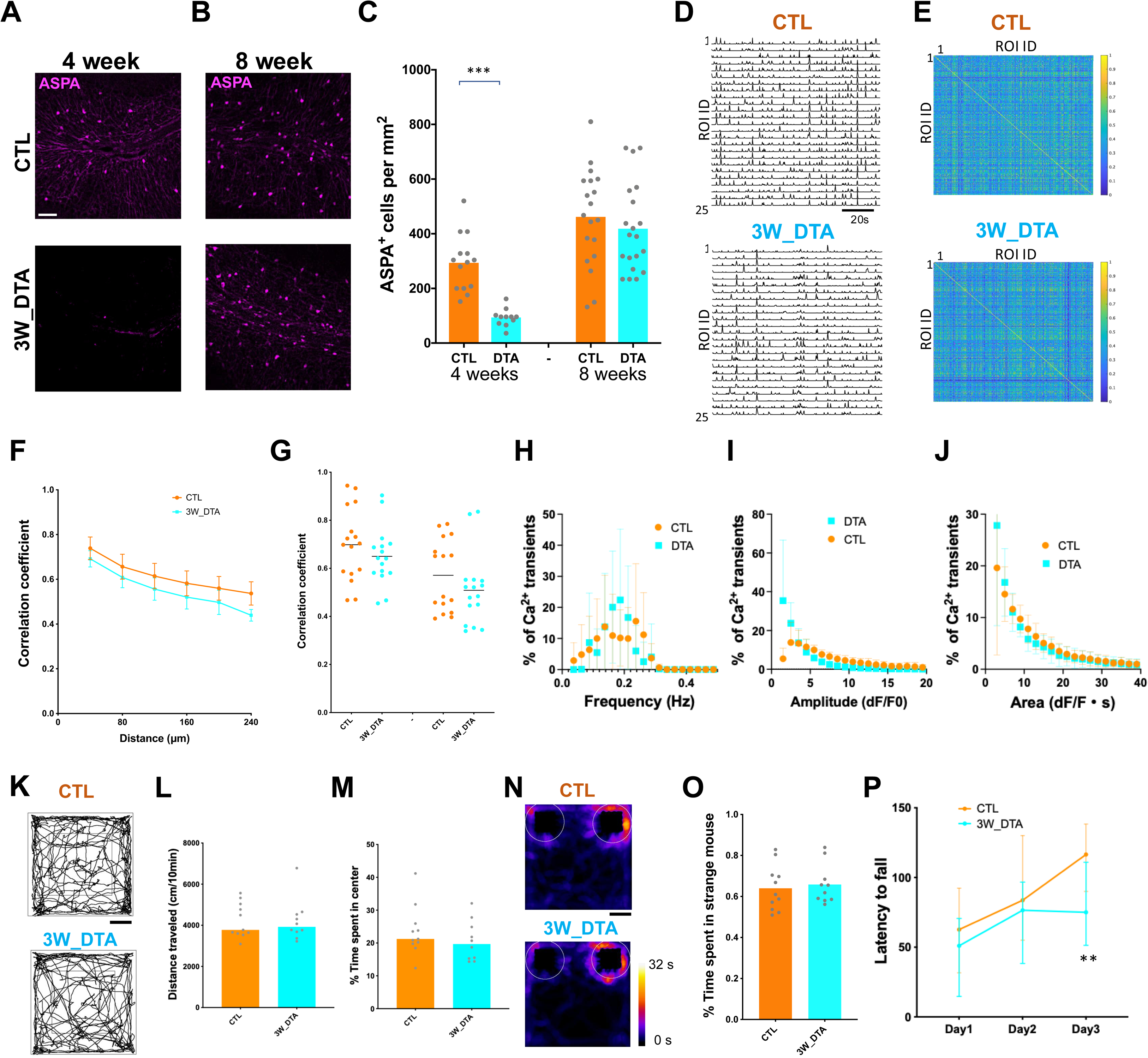
Oligodendrocyte reduction after 3 weeks old in mice does not affect PC activity synchrony and mouse behaviors except for motor learning. **A, B,** Fluorescence microscopy images comparing ASPA expression in the cerebellum of control (CTL) and 3-week DTA-treated (3W_DTA) mice at 4 weeks (**A**) and 8 weeks (**B**) old. ASPA-positive oligodendrocytes appear in pink. Scale bar, 50µm. **C,** Quantification of the number of ASPA-positive cells per mm² in the cerebellum of CTL and 3W_DTA mice at 4 and 8 weeks old, with data points for each field of view overlaid on the bar graphs (mean, 4 mice for each group at 4 weeks, 5 mice for each group at 8 weeks). *** p < 0.001 (Nested t-test). **D,** Representative calcium imaging traces from PCs in the cerebellar cortex of control (CTL) and 3-week DTA injected (3W_DTA) mice at 4weeks old, showing 25 regions of interest (ROIs). **E,** Correlation coefficient matrices of calcium transients between ROIs from CTL and 3W_DTA mice at 4 weeks old, indicating the level of synchrony, with warmer colors representing higher correlation coefficients. **F,** Line graph depicting mean correlation coefficients with increasing inter-ROI distance for CTL and 3W_DTA mice at 4 weeks old. Error bars indicate SEM. No significant group differences were observed (two-way ANOVA, 5 mice for each group). **G,** Scatter plot comparing the correlation coefficients of calcium transients in PCs between adjacent ROIs in CTL and 3W_DTA mice. No significant group differences were observed (Nested t-test, 5 mice for each). **H-J,** Frequency distributions of Frequency (H), Amplitude (I), and Area (J) of calcium transients in CTL (AAV-hMAG-mCherry) and 3W_DTA (AAV-hMAG-DTA) mice at 4 weeks. Circles and squares indicate data from individual ROIs, and bars represent SEM. There was no significant difference between groups (two-way ANOVA). **K-M,** Assessment of exploratory behavior and locomotor activity in control (CTL, orange, 11 mice) and 3 weeks DTA-injected (3W_DTA, cyan, 10 mice) mice in a novel environment, with trajectories of mouse exploratory behavior (**K**), distance traveled (**L**), and time spent in center (**M**) at P60-80. Bars and dots indicate median and data from individual mice, respectively. No significant group differences were observed (Mann-Whitney U tests). **N, O**, Representative images and quantitative analysis of sociality in CTL and 3W_DTA mice. A stranger mouse is present in the right cage and a doll is in the left cage. **N** displays the heatmap of mouse exploratory behavior, and (**O**) shows the percentage of time spent around strange mouse at P60-80 (11 mice for control, 10 mice for 3W_DTA). Bars and dots indicate median and data from individual mice, respectively. No significant group differences were observed (Mann-Whitney U tests). **P,** Line graph showing the median of latency to fall in the rotarod test over a three-day period for CTL and 3W_DTA mice at P60-80. Error bars indicate 95 % CI. Statistical significance is indicated by asterisks (**p < 0.01, Mann-Whitney U tests).

### Synchronized spontaneous activity during development regulates brain functions

To test the causal link between synchronized PC activity during developmental stage and cerebellar function, we used an experimental strategy that selectively manipulated CF activity, which drives PC calcium transients. We reduced synchronized spontaneous activity of PC population by expressing Kir2.1, a potassium channel that reduces neuronal excitability, in a subset of CFs projecting to PCs. Co-injection of AAVs with EGFP or Kir2.1-T2A-GFP under the control of TRE promoter and AAVs with tetracycline transactivator (tTA) under the control of Htr5b promoter (Dorgans *et al*., 2022) into the inferior olive at P7 labeled a subset of CFs in the cerebellum at P22 (Figure 7A-C). We quantified the proportion of PCs that received labeled CFs and found that PCs with GFP-labeled CFs were detected in 19.3 ± 11.9 % (mean ± S.D.) for control and 26.2 ± 11.8 % (mean ± S.D.) for Kir2.1 groups (Figure 7D). In vivo calcium imaging of CFs showed that activity of CFs with Kir2.1-T2A-jGCaMP7 was lower than that for CFs with only jGCaMP7 at P13-15 (Figure supplement 4A-D). In vivo calcium imaging of PC populations revealed that synchrony among distal PCs was unchanged, whereas synchrony among proximal PCs receiving Kir2.1-expressing CF inputs was significantly reduced at P13–15 compared with controls (Figure 7E-H). The frequency, amplitude and area of calcium transients were not significantly altered by Kir2.1 expression in a subset of CFs (Figure 7I-K). These results indicate that kir2.1 expression in a subset of CFs leads to reduced synchrony of proximal PC population activity.

**Figure 7.**
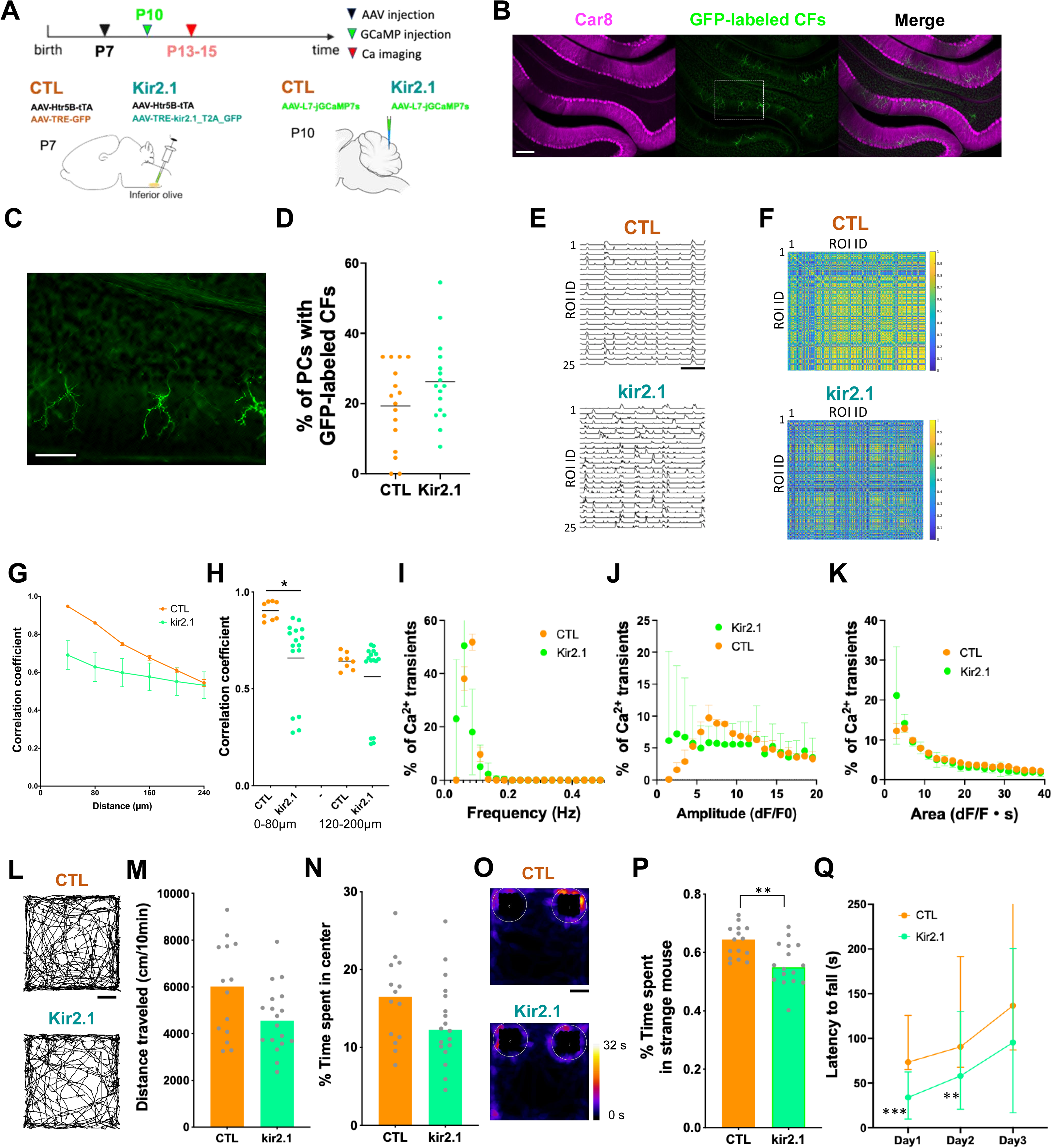
Kir2.1 expression in CFs reduces the synchrony of PC population spontaneous activity and leads to abnormal behaviors similar to the phenotype of mice with developmental oligodendrocyte reduction. **A**, Experimental scheme. AAV-Htr5B-tTA & AAV-TRE-GFP or AAV-Htr5b-tTA & AAV-Tre-kir2.1_T2A_GFP were injected into the cerebellum at P7, and AAV-L7-jGCaMP7f was injected into the inferior olive at P10 to label CFs. CF activity was imaged in the cerebellar cortex at P13-15. **B**, Fluorescent images of sagittal cerebellar sections showing Car8 immunoreactivity (Purkinje cells, magenta) and GFP expression (Climbing fibers (CFs), green). Scale bar, 200 µm. **C,** Higher magnification of the boxed area in **A** showing detailed GFP-labeled CF morphology. Scale bar, 100 µm. **D,** Scatter plot graph depicting the percentage of PCs with GFP-positive CFs among PC populations in CTL and Kir2.1 mice (17 sections from 10 mice for CTL and ). **E,** Representative time-course of spontaneous calcium transients from regions of interest (ROIs) in the mouse cerebellum at P13 showing spontaneous PC activity in control mice (CTL, top) and mice expressing Kir2.1 in a subset of CFs (kir2.1, bottom). Scale bar, 20 s. Y-axis is ΔF/F0 = (F - F0)/(F0). **F,** Correlation matrices for spontaneous PC activity between ROIs in control (top) and kir2.1-expressing (bottom) mice at P13. The color scale indicates correlation strength. **G**, Graph displaying the correlation coefficient of Ca^2+^ transients among PCs as a function of distance between adjacent PCs in CTL and Kir2.1 groups at P13-15 (4 mice for CTL, 8 mice for Kir2.1). Data points indicate the mean ± SEM. **H,** Scatter plot comparing correlation coefficients for ROI pairs within 0-80μm and 120-200μm separation distance ranges in CTL (orange) and kir2.1-expressing (cyan) mice at P13-15, with lines representing average values. Statistical significance is indicated by asterisks (*p < 0.05, Nested t-test, 4 mice for CTL, 8 mice for Kir2.1). **I-K**, Frequency distributions of Frequency (I), Amplitude (J), and Area (K) of calcium transients in CTL and Kir2.1 mice at P13-15. Circles and squares indicate data from individual ROIs, and bars represent SEM. There was no significant difference between groups (two-way ANOVA). **L-N,** Analysis of exploratory behavior and locomotor activity in CTL (orange) and Kir2.1-expressing (green) mice at P60-80 with distance traveled (**L, M**) and percentage of time spent in the center (**N**) in an open-field test (14 mice for CTL, 18 mice for Kir2.1). Bars and dots indicate median and data from individual mice, respectively. **O, P,** Representative images and quantitative analysis of sociality in CTL and Kir2.1-expressing mice at P62-80. (**O**) displays the heatmap of mouse exploratory behavior, and (**P**) shows the percentage of time spent around strange mouse. Bars and dots indicate median and data from individual mice, respectively. **Q**, Rotarod test. The line graph shows the median latency to fall across three days for CTL and Kir2.1-expressing mice at P64-80. Error bars indicate 95 % CI. Statistical significance is indicated by asterisks (**p < 0.01, *** p < 0.001, Mann-Whitney U tests).

Because Kir2.1 expression reduced PC synchrony during development, we next asked whether this manipulation also affected adult behavior. Behavioral testing at 2 months revealed that Kir2.1-expressing mice (injection of AAVs with Kir2.1 at P7) showed a trend toward decreased distance travelled and percentage time spent in center (Figure 7L-N). Kir2.1-expressing mice also exhibited reduced social interaction and impaired motor coordination, as shown by reduced social approach to a novel mouse and shorter latency to fall in the Rotarod test, respectively (Figure 7O-Q). These results indicate that the behavioral consequences of developmental oligodendrocyte ablation are phenocopied by experimentally reducing synchrony during development.

To directly test whether restoring PC synchrony in adulthood can rescue behavior, we expressed a red-shifted opsin (AAV-L7-rsChRmine) selectively in PCs of control and DTA mice and applied 590-nm light pulses (10 ms, 1 Hz) to the cerebellar vermis during behavioral testing (Figure 8A–B). Periodic optogenetic re-synchronization did not alter locomotor activity and anxiety-like behavior, as distance traveled and open-field center time remained low in DTA mice (Figure 8C-D). Unexpectedly, in control mice the same stimulation reduced center time (Figure 8E), indicating that artificially elevating activity synchrony and firing rate of PCs can enhance anxiety-like responses. In contrast, motor and social deficits in DTA mice were significantly improved. Optogenetic stimulation rescued sociability, as measured by the time spent interacting with a novel mouse (Figure 8F–G). The same stimulation restored rotarod performance to control levels (Fig. 8H). On day 1 without light stimulation, DTA mice with rsChrrmine expression performed similarly to DTA mice without rsChrmine and showed reduced latency to fall compared with control mice with rsChrmine. On day 2 and day 3 with light stimulation, DTA mice with rsChrmine showed longer latency to fall than DTA mice without rsChrmine, reaching values comparable to controls. These results demonstrate that reduced PC synchrony is the primary driver of the observed motor and social impairments, whereas anxiety-like behavior likely involves additional mechanisms. Together, these findings establish a causal link between PC synchrony and specific behavioral domains and show that periodic re-synchronization can normalize motor and social functions in adult mice that experienced developmental oligodendrocyte loss.

**Figure 8.**
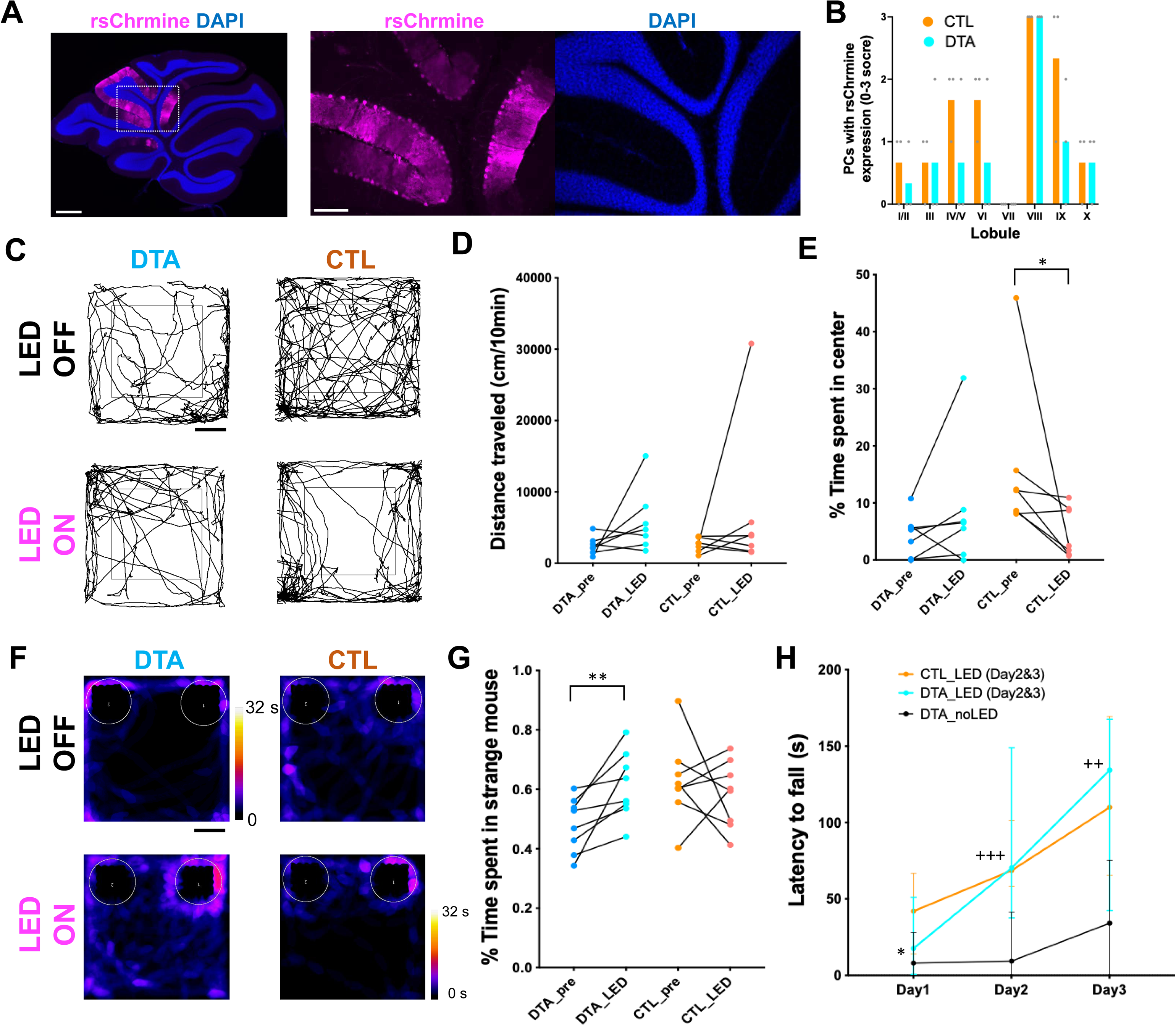
Optogenetic re-synchronization of PC activity rescues social and motor deficits but not anxiety-like behavior in adult mice with developmental oligodendrocyte ablation. **A,** Representative fluorescence images of cerebellar sagittal sections showing rsChrmine expression (magenta) in PCs and nuclear counterstain (DAPI, blue). Middle and right are the magnified images of dot region in left. Scale bars, 500 µm (left), 200 µm (middle, right). **B,** Bar graphs showing the distribution of PCs expressing rsChrmine in individual cerebellar lobules of control (CTL, orange, 3 mice) and DTA (cyan, 3 mice) mice. The number of rsChrmine-positive PCs in each lobule was evaluated using a semi-quantitative 4-point scale (0 = none, 1 = few, 2 = moderate, 3 = abundant). Data are presented as mean values across lobules, with individual dots indicating scores from each animal. **C,** Representative trajectories of exploratory behavior in the open-field test from DTA and control (CTL) mice with or without LED stimulation at P60. **D,** Quantification of total distance traveled in the open-field at P60-65. Each dot represents an individual mouse (7 mice for each). **E,** Quantification of percentage time spent in the center of the open-field. LED stimulation did not alter anxiety-like behavior in DTA mice, whereas in CTL mice it reduced center time, indicating increased anxiety-like responses. **F,** Representative heat maps of social exploration during the sociality test in DTA and CTL mice at P64 with or without LED stimulation. **G,** Quantification of sociability, measured as the percentage of time spent around a novel mouse at P64-68 (8 mice for each). Optogenetic stimulation rescued sociability in DTA mice to control levels. **H,** Rotarod test performance across three days at P66-70 (9 mice for CTL-LED, 8 mice for DTA-noLED, 8 mice for DTA-LED). Optogenetic stimulation restored motor coordination in DTA mice to control levels. Data represent median ± 95% CI. Statistical significance is indicated by asterisks (*p < 0.05, **p < 0.01, ***p < 0.001, ^++^p < 0.01, ^+++^p < 0.001; Mann–Whitney U test). + shows the significant difference between DTA_noLED and DTA_LED.

## Discussion

This study provides compelling evidence that oligodendrocytes play a crucial role beyond their traditional functions in myelination, extending to the regulation of synchronized spontaneous activity, and emphasizing the complexity of glial-neuronal interactions in the developing brain. Moreover, our research illuminates the significance of brain development following a proper course in brain function integrity

The interpretation of our synchrony measurements depends critically on the cellular sources of PC calcium signals. Our calcium imaging approach was designed to capture dendritic transients in PCs, which are overwhelmingly driven by CF complex spikes (Good *et al*., 2017). Previous work has shown that simple-spike (SS) bursts can evoke modest Ca²⁺ rises, but these are restricted to the soma and proximal dendrites of adult PCs and do not propagate into the distal dendritic arbor where our regions of interest were defined (Ramirez & Stell, 2016). Consistent with these, in vivo imaging showed that oligodendrocyte ablation reduced CF–CF synchrony at P14, directly linking oligodendrocytes to impaired CF coordination and PC hyposynchrony. While SS-dependent Ca²⁺ signals may still influence long-term circuit function, they likely do not contribute to the synchrony observed here. Future somatic imaging and electrophysiology will clarify whether oligodendrocytes also modulate SS activity.

In adult brain, adaptive myelination can induce and maintains synchrony of spontaneous activity among neural populations (Kato *et al*., 2020; Bonetto *et al*., 2021; Pajevic *et al*., 2023). Developmental oligodendrocytes have been also speculated to regulate the synchrony of neuronal activities (Jang *et al*., 2019), and our study now provides direct evidence for this role. Oligodendrocytes regulate synchrony in part through myelin formation and plasticity, which modulate axonal conduction velocity, while reciprocal neuronal signals drive myelination (Fields, 2015). Thus, bidirectional oligodendrocyte–neuron interactions can shape the timing of circuit activity, and alterations in these interactions may lead to desynchronization. Mechanistically, oligodendrocytes can regulate synchrony through at least two complementary routes. First, multilamellar myelin shortens membrane time constants and accelerates conduction velocity, thereby reducing temporal dispersion of CF inputs and sharpening coincidence detection in PCs. Second, mature oligodendrocytes provide metabolic support—including lactate, cholesterol, and mitochondrial intermediates—that sustains high-frequency axonal firing. Loss of this trophic support can prolong refractory periods and promote excessive desynchronization, even in the absence of overt demyelination.

Our findings reveal that the timing of PC desynchronization coincides with the initial appearance of oligodendrocytes and myelin in the cerebellum, with synchrony declining from P3–P5 (Good *et al*., 2017), just as myelin sheaths begin to emerge (Figure supplement Fig. 1A-B). However, developmental ablation of oligodendrocytes and myelin from P7–P21 reduced PC synchrony rather than precipitating its decline, arguing that early desynchronization is not triggered by myelin itself. Instead, our data point to non-myelin mechanisms—such as transient gap-junction coupling or shared climbing-fiber drive—as the agents of the initial fall in synchrony, with myelination coming online later to counterbalance these forces and stabilize the mature, low-synchrony regime. Consistent with this interpretation, in vivo imaging showed that oligodendrocyte loss specifically weakened climbing-fiber coordination at P14, directly linking developmental oligodendrocytes to the CF-driven component of PC synchrony. On this basis, we propose a two-phase model. Phase I (P3–P7): high early synchrony is reduced by non-myelin processes. Phase II (P7 onward): as oligodendrocytes proliferate and ensheath axons, they tune conduction velocity and provide metabolic support, thereby consolidating circuit timing and maintaining the adult low-synchrony regime.

Recent work further refines this framework by showing that single oligodendrocytes exhibit axonal selectivity that is set during development: in cerebellar white matter, roughly half of oligodendrocytes in adults preferentially myelinate PC axons, and this preference is markedly stronger at the onset of myelination; early-born oligodendrocytes preferentially ensheath PC axons in mature white matter, whereas later-differentiating oligodendrocytes predominantly myelinate non-PC axons (Battulga *et al*., 2025). These results imply that Phase II is not merely “more myelin,” but the emergence of cell-type–specific conduction control established within a critical developmental window. In our model, the selective, time-staggered ensheathment of PC versus non-PC axons narrows pathway-specific conduction dispersion and supports sustained firing, thereby preserving the low-synchrony state once it has been achieved. This selectivity also provides a mechanistic account for why oligodendrocyte ablation after P7 diminishes synchrony. Eliminating the early-born oligodendrocytes that normally establish preferential myelination of PC axons would remove the selective speeding and stabilization of PC conduction. Without this bias, action potential arrival times across PCs would become more dispersed, broadening conduction timing and weakening the temporal precision of climbing-fiber inputs. Because PCs project to the deep cerebellar nuclei (DCN), which in turn provide feedback to the inferior olive (IO)—the origin of CFs—disrupted PC synchrony would propagate through this loop, altering IO activity patterns and further degrading CF–PC coordination. Thus, even though the initial P3–P5 desynchronization is generated by non-myelin mechanisms, the later contribution of oligodendrocytes is essential to stabilize the entire PC–DCN–IO circuit and maintain the mature low-synchrony state.

Correlated spontaneous activity has been shown to play a pivotal role in neural development and the establishment of functional neural circuits especially in the developing visual system (Kirkby *et al*., 2013; Ackman & Crair, 2014). Retinal waves observed in various species, play a fundamental role in developing interconnections within the visual system including topographic maps. Recent study demonstrates that retinal wave is not merely random but possesses an intrinsic directionality that prefig the optic flow patterns encountered after eye opening. However, the effects of correlated spontaneous activity during development on adult brain function remain largely unknown. Our study suggests that synchronized spontaneous activity during development play a crucial role in the subsequent brain function. These studies provide new insights into specific information content carried by correlated spontaneous activity and their influence on the development of brain functions including higher-order function, extending beyond the previously understood role of these activity in neural circuit maturation.

Our findings demonstrate that the impact of oligodendrocyte loss on cerebellar synchronized activity and behavior is not merely transient but is gated by a narrow developmental window. Ablating oligodendrocytes during the second postnatal week—when myelination is beginning—produced a reduction in PC synchrony that was already evident at P14 and persisted unchanged into adulthood (P60–P70), even though oligodendrocyte density and MBP-positive myelin had fully recovered. The same early manipulation also led to lasting anxiety-like behavior, impaired motor coordination, and reduced sociability. By contrast, an equally robust ablation initiated after the third postnatal week (from P21) caused only a transient fall in oligodendrocyte numbers and had no measurable effect on synchrony or behavior. These results indicate that developmental myelination and/or the associated metabolic support provided by oligodendrocytes is indispensable for locking in the low-synchrony, high-fidelity state of mature cerebellar circuits, whereas the disruption outside this critical period confers little impact. Thus, the lasting network and behavioral consequences we observe stem from a time-limited developmental requirement for oligodendrocyte function rather than from chronic hypomyelination per se.

There is growing evidence that oligodendrocytes and myelin affect learning, memory, and cognitive function (de Faria *et al*., 2021; Munyeshyaka & Fields, 2022). Moreover, numerous studies have reported the changes of white matter, oligodendrocyte, and myelin in schizophrenia, autism spectrum disorders, depression, and anxiety, suggesting that the deficit of oligodendrocytes and myelin may underlie multiple brain disorders (Edgar & Sibille, 2012; de Faria *et al*., 2021). However, it is unclear when, where, and how oligodendrocytes and myelin are involved in the functionalization of brain and the pathogenesis of these brain disorders. Our data showing that synchronized spontaneous activity during development, controlled by oligodendrocytes, is a precursor to the maturation of functional neural circuits and brain functions provides a novel lens through which to examine the etiology of disorders characterized by synaptic imbalances and dysregulation of neural activity. Supporting this view, we found that reduced PC synchrony persists into adulthood after developmental oligodendrocyte loss, and that optogenetic restoration of synchrony rescues social and motor behaviors. Importantly, this effect cannot be explained by a mere increase in firing frequency. PC calcium event rates did not differ between control and DTA mice, and optogenetic stimulation of control mice did not further enhance social interaction or rotarod performance. Thus, oligodendrocyte and myelination abnormalities, and the resulting disruption of synchronized activity in neuronal populations, may be new therapeutic targets for multiple brain disorders.

Mounting evidence indicates that even subtle shifts in the temporal coordination of PC ensembles can cascade through cerebello–thalamo–cortical pathways and reshape distributed brain dynamics underlying motor planning, emotional regulation, and social cognition (Guell *et al*., 2018; Stoodley & Schmahmann, 2018; Lindeman *et al*., 2021; McAfee *et al*., 2021). Complex-spike synchrony is organized by aldolase-C module boundaries, generating modular timing channels within the cerebellar cortex (Tsutsumi *et al*., 2015), and disrupting synchrony in a single module is sufficient to alter behavior in real time (Tsutsumi *et al*., 2019). During reinforcement learning, these modules reorganize their synchrony into a handful of low-dimensional components that track motor initiation, error prediction, and reward valuation (Hoang *et al*., 2023), highlighting flexible coupling between cerebellar timing and higher-order cognitive circuits. To test the behavioral relevance of this persistent hyposynchrony we used optogenetics to transiently resynchronize PC ensembles in adult mice with DTA during development. This manipulation restored sociability and motor coordination but did not normalize anxiety-like behavior. These findings indicate that different behavioral domains vary in their dependence on cerebellar synchrony, with motor and social functions strongly reliant on precise Purkinje cell activity timing while anxiety involves additional circuit mechanisms. More broadly, neural synchrony does not operate in isolation within a single structure but propagates through cerebello–thalamo–cortical loops to influence prefrontal, limbic and parietal networks. Thus, even localized perturbations in Purkinje synchrony can acquire system-level significance and shape distributed brain states that support cognition, affect and social interaction.

Given the pivotal role of oligodendrocytes in neural circuit maturation, future research should aim to further dissect the molecular and cellular mechanisms underpinning their influence on synchronized spontaneous activity. Understanding the signaling pathways and interaction networks involved could open new therapeutic avenues for addressing developmental brain disorders and enhancing neural repair after injury.

## Acknowledgments

We thank Dr. Daisuke Tanaka, Dr. Shizuki Inaba, Dr. Ayaka Nakai, and other lab members of Uesaka lab for helpful comments and discussions.

## Funding

This work is supported by MEXT Grants-in-Aid for Scientific Research on transformative research area B “Design-build of brain through spontaneous brain multimodal activity” (KAKENHI 22H05092, 22H05093), the Asahi Glass Foundation, Toray Foundation, Takeda Science Foundation, Tokumori Yasumoto Memorial Trust, and Uehara Memorial Foundation to N.U.

## Author contributions

Initiation and conceptualization: N.U. Investigation: R.M., K.G., M.S., and N.U. Writing - original draft: N.U. Writing - editing: R.M., K.G., N.U., with inputs from all authors. Visualization: R.M., and N.U. Supervision: N.U. Funding acquisition: N.U.

## Competing interests

The authors declare no conflicts of interest.

## Data and materials availability

All data are available in the main text or the supplementary materials.

## Materials availability

All unique/stable reagents generated in this study are available from the lead contact upon request with a completed Materials Transfer Agreement.

## Data and code availability

Requests for behavioral and/or calcium imaging data reported in this paper will be shared by the lead contact upon reasonable request.

Any additional information required to reanalyze the data reported in this paper is available from the lead contact upon reasonable request.

## Supporting Information

**Fig. S1.**
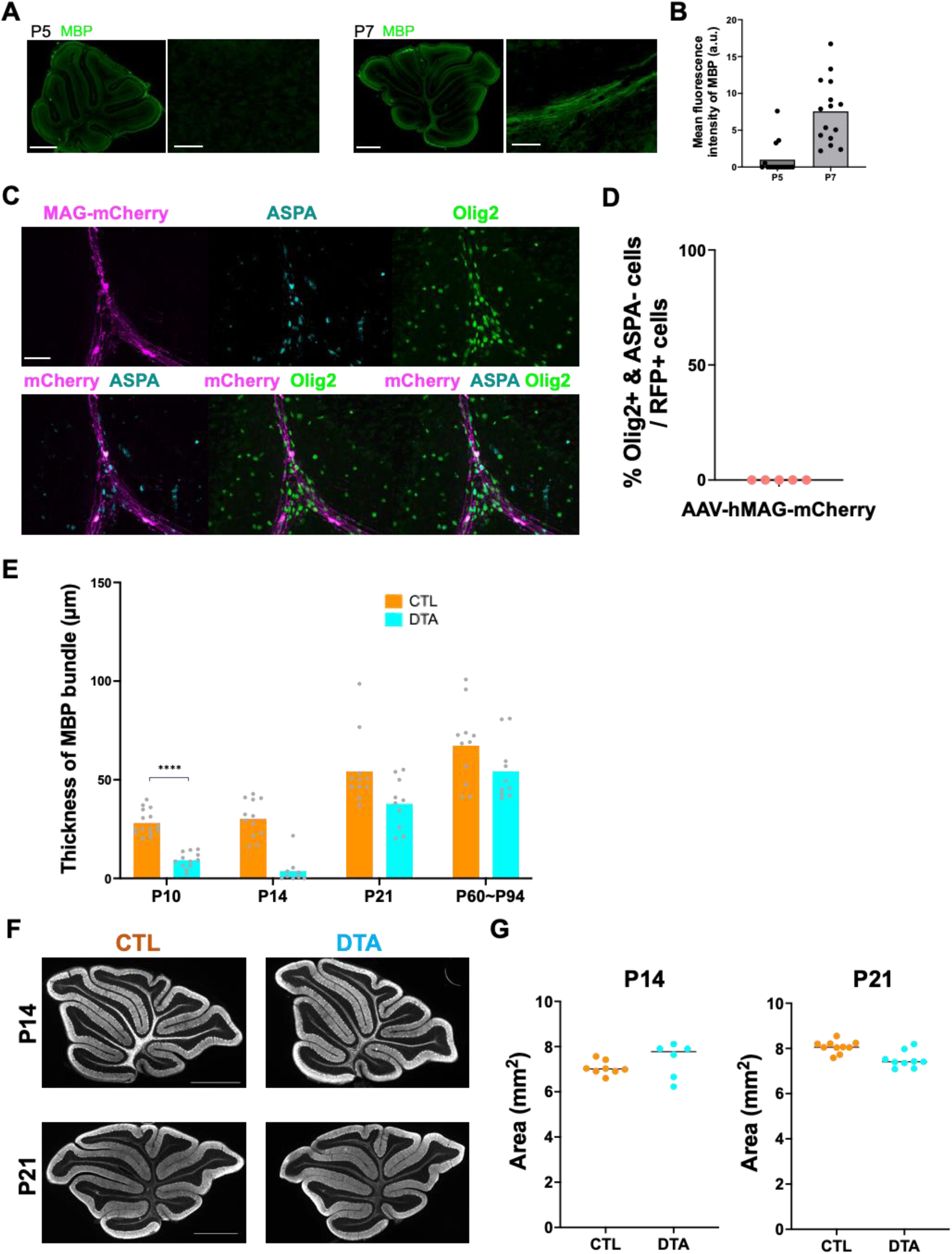
Targeted myelin disruption by oligodendrocyte-specific AAVs. **A**, Representative images of MBP immunostaining in sagittal sections of control mice at P5 and P7, shown at low magnification (left) and higher magnification of the white matter (right). Scale bars, 500 µm (low magnification), 50 µm (high magnification). **B**, Bar graphs showing the mean fluorescence intensity of MBP at P5 and P7, derived from three mice in each group. **C**, Representative immunofluorescence images showing the specificity of MAG-mCherry expression at P14. mCherry (magenta) co-localizes with ASPA-positive oligodendrocytes (cyan) and Olig2-positive oligodendrocyte lineage cells (green). Notably, mCherry was absent in ASPA-negative, Olig2-positive immature oligodendrocytes. Scale bar, 50 µm. **D**, Quantification of the proportion of mCherry+ cells that were ASPA- and Olig2+ cells in total mCherry+ cells (5 mice). Dots represent individual mice. **E**, Quantification of the thickness of MBP bundle in CTL and DTA mice at P10, P14, P21, and adulthood (P60–P94). The thickness was significantly reduced at P10 and tended to be reduced at P14 (p = 0.13) but recovered to control levels by P21 and adulthood (n = 4 mice per group). Bars indicate mean, dots represent individual fields of view. ****p < 0.0001 (nested t-test). **F**, Low-magnification images of parvalbumin-stained sagittal sections showing overall cerebellar morphology and foliation in CTL and DTA mice at P14 and P21. No gross abnormalities were observed. Scale bar: 1 mm. **G**, Quantification of total cerebellar area measured from sagittal sections at P14 (n = 8 mice for control, n = 6 mice for DTA) and P21 (n = 10 mice for control, n = 9 mice for DTA). No significant differences were found between CTL and DTA mice (Mann–Whitney U test). Bars indicate median, dots represent the data for individual animals.

**Fig. S2.**
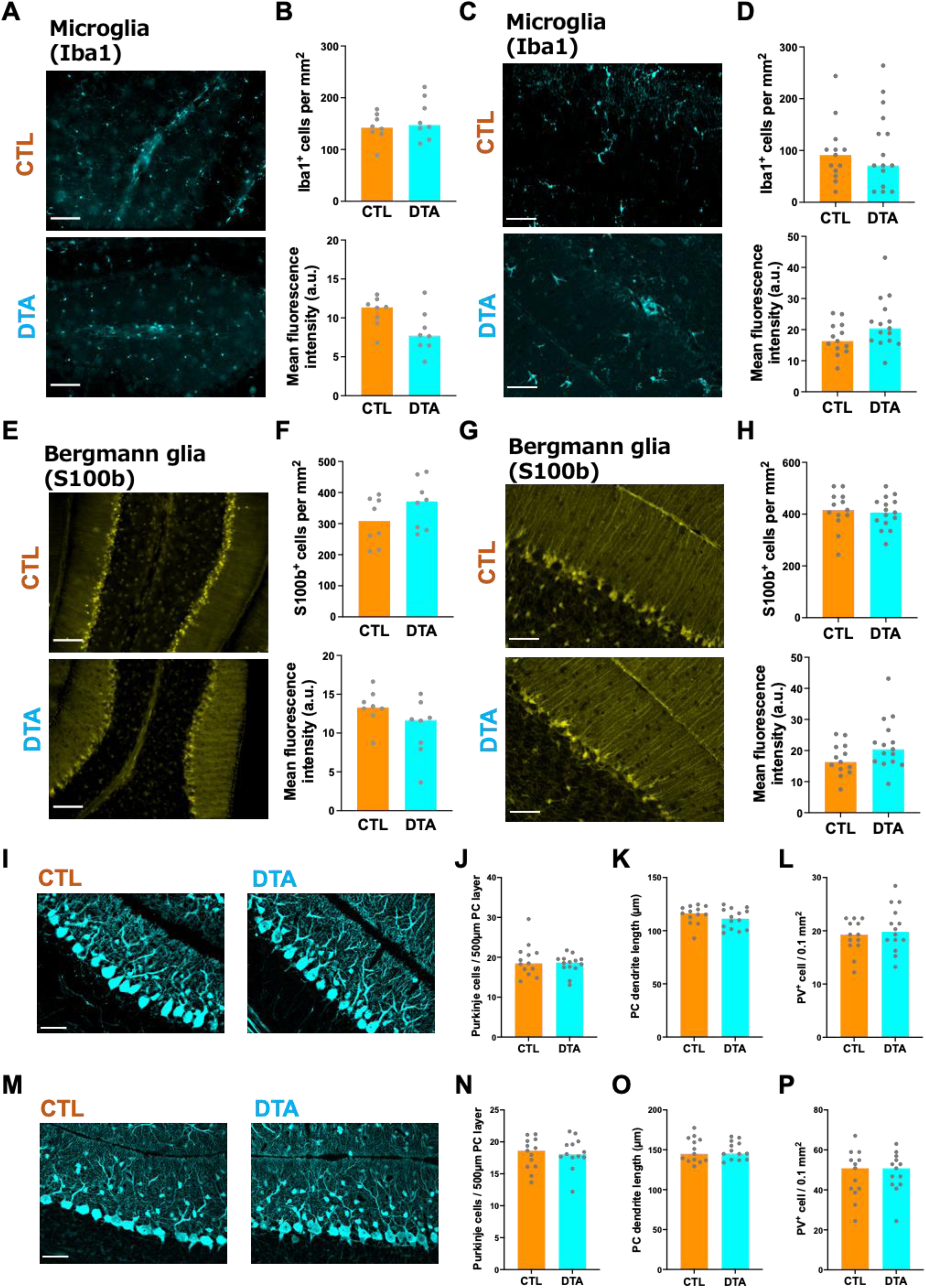
Oligodendrocyte ablation does not alter other glial or neuronal populations in the developing cerebellum. **A, B,** Representative images at P14 and quantification of Iba1 immunostaining at P13-15 showing microglial density and mean fluorescence intensity in control (CTL) and DTA mice (8 mice for each). Scale bar, 50 µm. Bars indicate median, dots represent the data for individual animals. No significant differences were found between CTL and DTA mice (Mann–Whitney U test). **C, D,** Representative images at P21 and quantification of Iba1 immunostaining at P21-28. Bars indicate median, dots represent the data for individual animals. No significant differences were observed between CTL and DTA groups (13 mice for CTL, 15 mice for DTA, Mann–Whitney U test). **E, F,** Representative images at P14 and quantification of S100β immunostaining at P13-15 showing Bergmann glial density and mean fluorescence intensity (9 mice for CTL, 8 mice for DTA). Scale bar, 50 µm. Bars indicate median, dots represent the data for individual animals. No significant differences were found between CTL and DTA mice (Mann–Whitney U test). **G, H,** Representative images at P21 and quantification of S100β immunostaining at P21-28 (13 mice for CTL, 15 mice for DTA). Scale bar, 50 µm. Bars indicate median, dots represent the data for individual animals. No significant differences were found between CTL and DTA mice (Mann–Whitney U test). **I–L,** Parvalbumin (PV) immunostaining at P14. Quantification includes PC number, PC dendritic arbor size, and molecular layer PV+ interneuron density (13 mice for CTL, 15 mice for DTA). Scale bar, 50 µm. Bars indicate median, dots represent the data for individual animals. No significant differences were found between CTL and DTA mice (Mann–Whitney U test). **M–P,** PV immunostaining at P21. Quantification includes PC number, PC dendritic arbor size, or PV+ interneuron density (13 mice for each). Scale bar, 50 µm. Bars indicate median, dots represent the data for individual animals. No significant differences were found between CTL and DTA mice (Mann–Whitney U test).

**Fig. S3.**
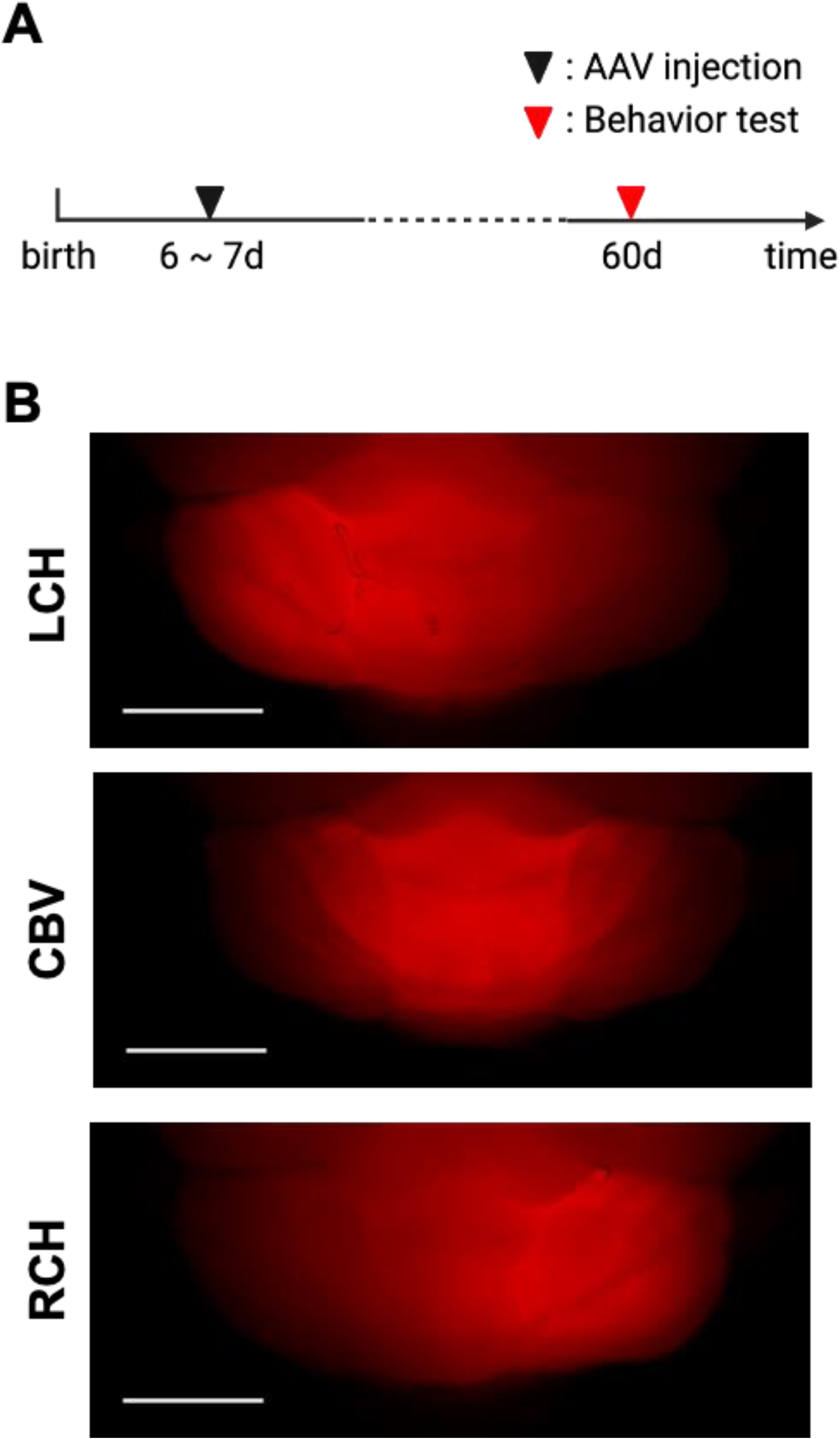
Timeline of AAV injection and behavioral assessment for mice with the injection into distinct cerebellar regions. **A,** Schematic timeline illustrating the experiments from birth to behavioral assessment, with AAV injections performed at P6–P7 and subsequent behavioral assessments conducted at around P60. **B,** Representative whole-cerebellum images of mCherry expression highlighting three different regions: left cerebellar hemisphere (LCH), cerebellar vermis (CBV), and right cerebellar hemisphere (RCH). Scale bars, 2 mm.

**Fig. S4.**
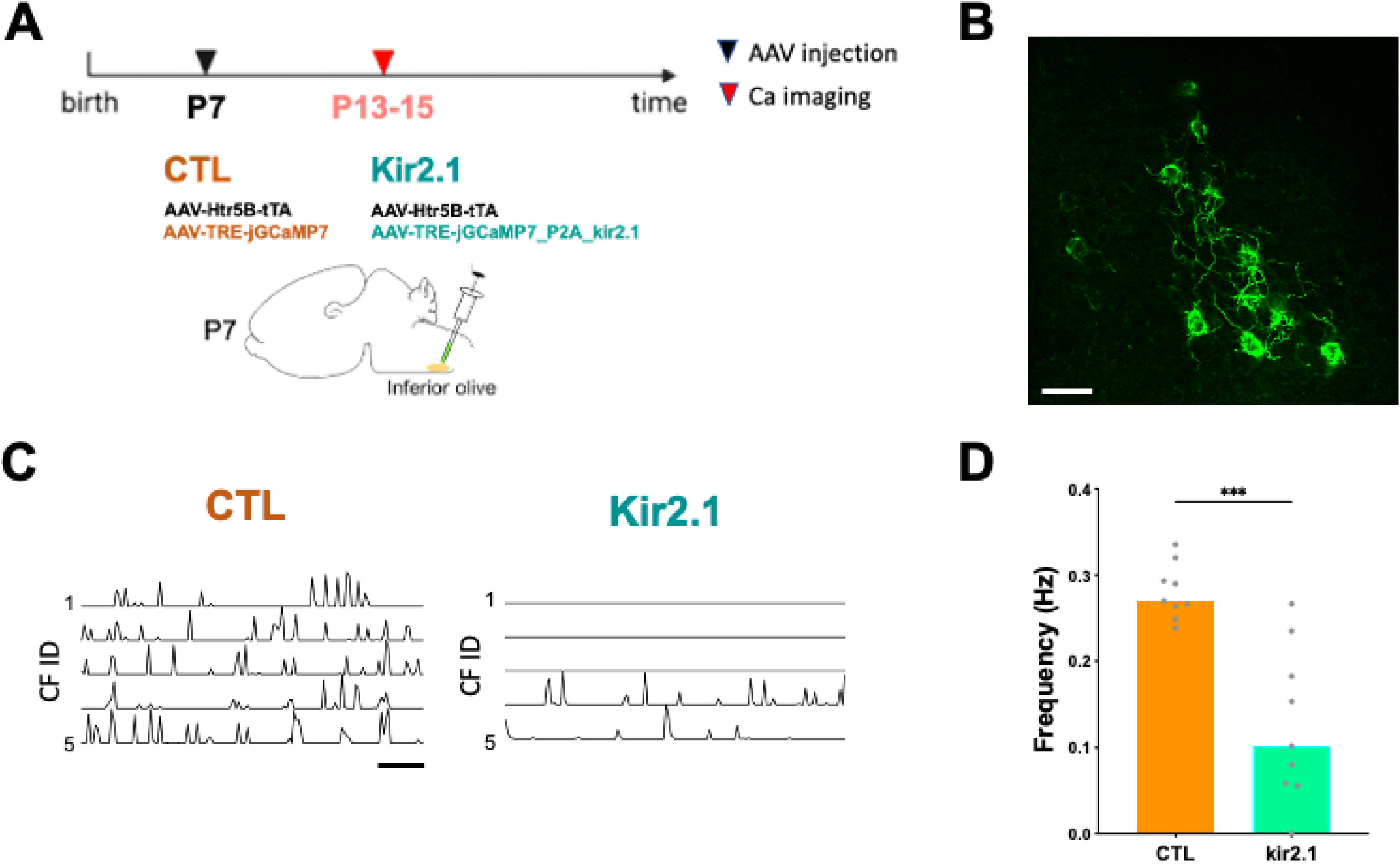
Kir2.1 expression in CFs reduces the CF activity. **A**, Schematic timeline of the experimental protocol. At P7, AAV injections are administered to either overexpress Kir2.1 and GFP for Kir2.1 or only GFP for control (CTL). Ca^2+^ imaging is conducted at P13-15. **B**, Representative image of jGCaMP7s-labeled CFs in the Kir2.1 experimental group, captured using two-photon laser scanning microscopy. Scale bar, 50 µm. **C**, Representative traces of calcium transients in CTL and Kir2.1-overexpressing CFs. Each tick mark represents a single transient. Scale bar, 20 s. **D**, Bar graph showing the median frequency of calcium transients in CTL and Kir2.1-overexpressing CFs, with data points of individual cells. ***p < 0.001 (Mann-Whitney U test).

## Notes

### Competing Interest Statement

The authors have declared no competing interest.

### Summary of Updates

Revise Figure7. Previous Figure 7 graphs are wrong. We have changed thesee graphs.

