## Supplementary_table1 for "Developmental oligodendrocytes regulate brain function through the mediation of synchronized spontaneous activity"

| | | Control_P10 | | | |
|---|---|---|---|---|---|
| mouse1 | mouse2 | mouse3 | mouse4 | mouse1 | mouse2 |
| 49.7588278 | 101.839348 | 135.219404 | 158.353081 | 0 | 69.9884793 |
| 72.8805652 | 12.1548619 | 129.645301 | 79.294675 | 31.8322077 | 0 |
| 96.618449 | 73.2810615 | 0 | 5.59151991 | 5.26049935 | 0 |
| 90.1256654 | 98.2400345 | 65.4586596 | 89.2528159 | 0 | 6.13347244 |
| 69.464485 | | 0 | 51.3091216 | 0 | 69.2053567 |
| | | 30.5276382 | | | |
| | | 83.7755955 | | | |

| DTA_P10 | | | | Control_P14 | |
|---|---|---|---|---|---|
| mouse3 | mouse4 | mouse1 | mouse2 | mouse3 | mouse4 |
| 4.70832903 | 49.9254276 | 157.605589 | 110.813697 | 177.674398 | 117.310345 |
| 0 | 45.8324019 | 152.129636 | 11.4933971 | 35.7874972 | 94.6753247 |
| 23.6734429 | 0 | 173.796584 | 135.649615 | 96.3465636 | 47.393617 |
| 0 | 0 | 171.31781 | 194.868695 | 198.608123 | 214.863158 |
| 37.2052669 | 70.6207164 | 49.646035 | | 10.2946993 | 65.6756757 |
| | 0 | | | 199.582133 | |
| | | | | 140.733591 | |

| | | DTA_P14 | | | |
|---|---|---|---|---|---|
| mouse1 | mouse2 | mouse3 | mouse4 | | mouse1 |
| 0 | 3.55516098 | 0 | 21.2948729 | | 339.444735 |
| 0 | 0 | 34.7522245 | 0 | | 329.886251 |
| 0 | 15.4657694 | 9.64388495 | 74.7266515 | | 222.80429 |
| 40.380545 | 41.9508996 | 21.8494268 | 91.6340782 | | 405.375348 |
| 0 | 100.423003 | 0 | 0 | | 229.95291 |
| | | | 0 | | |

| Control_P21 | | | | DTA_P21 | | |
|---|---|---|---|---|---|---|
| mouse2 | mouse3 | mouse4 | | mouse1 | mouse2 | mouse3 |
| 295.676206 | 426.191172 | | | 181.806569 | 318.787129 | 293.020493 |
| 147.220919 | 304.232909 | 277.233511 | | 238.5 | 134.289474 | 147.713084 |
| 293.410764 | 190.638075 | 267.085766 | | 269.232955 | 221.857776 | 306.138472 |
| 196.520034 | 214.331075 | 255.332254 | | 289.366295 | 271.763829 | 178.301887 |
| | 178.468695 | 305.054752 | | 208.027295 | 185.90194 | 191.195638 |
| | 332.525587 | 210.19246 | | | | |
| | 252.12614 | | | | | |

| mouse4 | | Control_P60-94 | | | |
|---|---|---|---|---|---|
| | | mouse1 | mouse2 | mouse3 | mouse4 |
| 210.224932 | | 488.952731 | 528.037383 | 549.759123 | 565.78009 |
| 242.876181 | | 399.687909 | 374.313008 | 557.580521 | 390.338454 |
| 368.980477 | | 512.202232 | 561.901266 | 637.970383 | 535.443427 |
| 236.115909 | | | | 510.05825 | 308.335378 |
| 293.183189 | | | | | |
| 46.6374942 | | | | | |

| | | DTA_P60-94 | |
|---|---|---|---|
| mouse1 | mouse2 | mouse3 | mouse4 |
| 472.088221 | 627.649857 | 489.017874 | 375.546374 |
| 438.550296 | 831.900698 | 465.98449 | 609.341449 |
| 800.797466 | 466.342939 | 694.272128 | 339.065671 |
| 877.388295 | 595.356755 | | |

| | | Control_P10 | | | | |
|---|---|---|---|---|---|---|
| mouse1 | mouse2 | mouse3 | mouse4 | | mouse1 | mouse2 |
| 3301.60078 | 4121.88 | 5853.52078 | 6915.84078 | | 71.52078 | 101.92 |
| 8583.83922 | 8717.08078 | 5321.4 | 5906.47922 | | 41.16 | 20.56078 |
| 1573.88 | 3899.43922 | 2700.88 | 7105 | | 118.5992 | 106.8392 |
| 5613.44 | 5963.28078 | 5908.40078 | | | | |

| DTA_P10 | | | Control_P14 | | | |
|---|---|---|---|---|---|---|
| mouse3 | mouse4 | | mouse1 | mouse2 | mouse3 | mouse4 |
| 168.56 | 338.0808 | | 4153.24 | 7611.67922 | 8048.72078 | 1629.75922 |
| 339.08 | 49.96078 | | 4675.56078 | 7055.03922 | 8442.71922 | 7461.72 |
| 662.48 | 3.92 | | 8117.32078 | 1662.08 | 4138.52078 | 6455.27922 |

| | | DTA_P14 | | | |
|---|---|---|---|---|---|
| mouse1 | mouse2 | mouse3 | mouse4 | | mouse1 |
| 0 | 19.6 | 3.92 | 0 | | 8960.15922 |
| 0 | 87.20078 | 1.96 | 1.96 | | 9179.64078 |
| | | | 20.56078 | | 9639.28 |

| | Control_P21 | | | | DTA_P21 |
| --- | --- | --- | --- | --- | --- |
| mouse2 | mouse3 | mouse4 | mouse1 | mouse2 | mouse3 |
| 9755.88078 | 9548.12078 | 8843.52 | 7778.241 | 9698.08 | 4929.4 |
| 9180.64 | 8347.64 | 9751 | 7883.12 | 9454.079 | 9643.2 |
| 9632.40078 | 9277.67922 | | 9397.201 | | |

| | | | | | |
|---|---|---|---|---|---|
| | | | | Control_P60-94 | |
| mouse4 | | mouse1 | mouse2 | mouse3 | mouse4 |
| 9492.28 | | 9717.68 | 9378.6 | 8907.20078 | 9602.04 |
| 1511.16 | | 9282.56 | 9102.24 | 9755.88078 | 9517.76 |
| 7034.44 | | 7668.48078 | 9460.92 | 9784.32 | |

| | | DTA_P60-94 | |
|---|---|---|---|
| mouse1 | mouse2 | mouse3 | mouse4 |
| 9330.599 | 8900.36 | 8988.56 | 8167.32 |
| 8962.081 | 9269.801 | 8460.359 | 9101.279 |
| | 8387.839 | | |

| | | Control_P10 | | | | |
|---|---|---|---|---|---|---|
| mouse1 | mouse2 | mouse3 | mouse4 | | mouse1 | mouse2 |
| 20.88 | 23.04 | 20.16 | 30.6 | | 4.32 | 11.88 |
| 39.96 | 37.08 | 23.04 | 31.32 | | 6.48 | 2.16 |
| 26.28 | 24.12 | 36 | 34.92 | | 8.64 | 14.4 |
| 25.2 | 24.84 | 25.2 | | | | |

| DTA_P10 | | | | | Control_P14 | |
|---|---|---|---|---|---|---|
| mouse3 | mouse4 | | mouse1 | mouse2 | mouse3 | mouse4 |
| 13.68 | 10.44 | | 21.6 | 32.4 | 42.84 | 16.92 |
| 14.76 | 8.64 | | 23.04 | 29.16 | 39.96 | 27.72 |
| 9.36 | 6.48 | | 40.68 | 41.04 | 16.2 | 31.68 |

| | | DTA_P14 | | |
|---|---|---|---|---|
| mouse1 | mouse2 | mouse3 | mouse4 | mouse1 |
| 0 | 19.6 | 1.96 | 1.96 | 8960.15922 |
| 0 | 87.20078 | 0 | 20.56078 | 9179.64078 |
| | 3.92 | | | 9639.28 |

| Control_P21 | | | DTA_P21 | | |
|---|---|---|---|---|---|
| mouse2 | mouse3 | mouse4 | mouse1 | mouse2 | mouse3 |
| 9755.88078 | 9548.12078 | 8843.52 | 7778.241 | 9698.08 | 9643.2 |
| 9180.64 | 8347.64 | 9751 | 7883.12 | 9454.079 | 9492.28 |
| 9632.40078 | 9277.67922 | | 9397.201 | 4929.4 | |

| | | Control_P60-94 | | | |
|---|---|---|---|---|---|
| mouse4 | | mouse1 | mouse2 | mouse3 | mouse4 |
| 1511.16 | | 9717.68 | 9378.6 | 8907.20078 | 9602.04 |
| 7034.44 | | 9282.56 | 9102.24 | 9755.88078 | 9517.76 |
| | | 7668.48078 | 9460.92 | 9784.32 | |

DTA_P60-94

| mouse1 | mouse2 | mouse3 | mouse4 |
|---|---|---|---|
| 9330.599 | 8900.36 | 8988.56 | 8167.32 |
| | 9269.801 | | |
| | 8387.839 | 8460.359 | 9101.279 |
| 8962.081 | | | |

| | | CTL 0-80μm | | | | |
|---|---|---|---|---|---|---|
| mouse1 | mouse2 | mouse3 | mouse4 | mouse5 | | mouse1 |
| 0.905193 | 0.741835 | 0.79329 | 0.811485 | 0.739528 | | 0.858098 |
| 0.858576 | 0.606863 | 0.683496 | 0.682442 | 0.61778 | | 0.743044 |
| 0.750626 | 0.69544 | 0.726591 | 0.695517 | 0.668368 | | 0.66689 |
| 0.648968 | 0.606714 | 0.648108 | 0.604331 | 0.561488 | | 0.584103 |

DTA 0-80μm

| mouse2 | mouse3 | mouse4 | mouse5 | mouse6 | mouse7 |
|---|---|---|---|---|---|
| 0.599792 | 0.528765 | 0.228595 | 0.526574 | 0.756936 | 0.671273 |
| 0.506506 | 0.475224 | 0.055729 | 0.220633 | 0.570846 | 0.502939 |
| 0.510614 | 0.295902 | 0.23275 | 0.481147 | 0.616576 | 0.574046 |
| 0.470126 | 0.254667 | 0.06955 | 0.228432 | 0.471902 | 0.454255 |

CTL 120-200µm

| mouse1 | mouse2 | mouse3 | mouse4 | mouse5 | | mouse1 |
|---|---|---|---|---|---|---|
| 0.702061 | 0.418138 | 0.578917 | 0.62568 | 0.644015 | | 0.573134 |
| 0.333927 | 0.654795 | 0.37874 | 0.548757 | 0.644037 | | 0.56964 |
| 0.249658 | 0.601684 | 0.485494 | 0.479775 | 0.294411 | | 0.483577 |

DTA 120-200µm

| mouse2 | mouse3 | mouse4 | mouse5 | mouse6 | mouse7 |
|---|---|---|---|---|---|
| 0.397446 | 0.378884 | -0.00244 | -0.04819 | 0.250555 | 0.315349 |
| 0.395566 | 0.394958 | -0.01241 | -0.03847 | 0.224936 | 0.300743 |
| 0.381985 | 0.18362 | -0.03407 | 0.040789 | 0.272283 | 0.305855 |

| | | CTL 0-80µm | | | | |
|---|---|---|---|---|---|---|
| mouse1 | mouse2 | mouse3 | mouse4 | mouse5 | mouse6 | mouse1 |
| 0.641609 | 0.614071 | 0.534597 | 0.516584 | 0.52269 | 0.492277 | 0.462047 |
| 0.584231 | 0.557234 | 0.532818 | 0.559615 | 0.565653 | 0.541039 | 0.262713 |
| 0.933327 | 0.890364 | 0.888721 | 0.864217 | 0.854016 | 0.846975 | 0.79603 |
| 0.734483 | 0.715809 | 0.713729 | 0.704843 | 0.690871 | 0.676688 | 0.46365 |

| | | DTA 0-80μm | | |
| --- | --- | --- | --- | --- |
| mouse2 | mouse3 | mouse4 | mouse5 | mouse6 |
| 0.424301 | 0.33458 | 0.294909 | 0.30186 | 0.279403 |
| 0.240235 | 0.258979 | 0.236113 | 0.198805 | 0.166278 |
| 0.639482 | 0.592262 | 0.58587 | 0.542117 | 0.520075 |
| 0.466333 | 0.433652 | 0.389543 | 0.426968 | 0.342971 |

CTL 120-200μm

| mouse1 | mouse2 | mouse3 | mouse4 | mouse5 | mouse6 | mouse1 |
|---|---|---|---|---|---|---|
| 0.466167 | 0.479619 | 0.469532 | 0.469514 | 0.462127 | 0.442091 | 0.137816 |
| 0.460475 | 0.451649 | 0.454705 | 0.448395 | 0.463161 | 0.417556 | 0.117225 |
| 0.623685 | 0.607393 | 0.607594 | 0.593217 | 0.592496 | 0.592438 | 0.255334 |
| 0.569948 | 0.553072 | 0.556885 | 0.551061 | 0.554295 | 0.538756 | 0.232234 |

DTA 120-200μm

| mouse2 | mouse3 | mouse4 | mouse5 | mouse6 |
| --- | --- | --- | --- | --- |
| 0.159631 | 0.107289 | 0.186545 | 0.143826 | 0.17469 |
| 0.136764 | 0.162031 | 0.135723 | 0.18995 | 0.112016 |
| 0.242996 | 0.310925 | 0.220053 | 0.196852 | 0.223832 |
| 0.202472 | 0.241492 | 0.267203 | 0.231295 | 0.206513 |

4 weeks

| | CTL | | |
|---|---|---|---|
| mouse1 | mouse2 | mouse3 | mouse4 |
| 208 | 304 | 200 | 328 |
| 200 | 152 | 304 | 280 |
| 408 | 296 | 408 | 176 |
| 328 | 520 | | |

8 weeks

| | | CTL | | |
|---|---|---|---|---|
| mouse1 | mouse2 | mouse3 | mouse4 | mouse5 |
| 150 | 384 | 468 | 630 | 378 |
| 540 | 264 | 810 | 600 | 132 |
| 660 | 510 | 552 | 318 | 594 |
| 444 | 450 | 300 | 594 | |

DTA

| mouse1 | mouse2 | mouse3 | mouse4 |
|---|---|---|---|
| 96 | 102 | 96 | 36 |
| 84 | 96 | 66 | 126 |
| 96 | 162 | 72 | |

DTA

| mouse1 | mouse2 | mouse3 | mouse4 | mouse5 |
|---|---|---|---|---|
| 558 | 396 | 516 | 336 | 420 |
| 234 | 438 | 258 | 318 | 714 |
| 234 | 270 | 318 | 570 | 714 |
| 312 | 234 | 390 | 702 | 438 |

| | | CTL 0-80μm | | | | |
|---|---|---|---|---|---|---|
| mouse1 | mouse2 | mouse3 | mouse4 | mouse5 | | mouse1 |
| 0.943926 | 0.752921 | 0.54854 | 0.594899 | 0.86707 | | 0.691512 |
| 0.724063 | 0.93324 | 0.73072 | 0.470223 | 0.58741 | | 0.580854 |
| 0.576988 | 0.70326 | 0.8775 | 0.69802 | 0.46747 | | 0.724532 |
| | | | | 0.68989 | | |

| | DTA 0-80µm | | |
|---|---|---|---|
| mouse2 | mouse3 | mouse4 | mouse5 |
| 0.687278 | 0.569525 | 0.466764 | 0.876906 |
| 0.903504 | 0.702276 | 0.64253 | 0.5814411 |
| 0.6693202 | 0.588892 | 0.618157 | 0.4543177 |
| | | | 0.6408947 |

CTL 120-200µm

| mouse1 | mouse2 | mouse3 | mouse4 | mouse5 |
|---|---|---|---|---|
| 0.784447 | 0.669022 | 0.40837 | 0.457852 | 0.73397 |
| 0.482071 | 0.7769 | 0.65359 | 0.396832 | 0.45291 |
| 0.415137 | 0.46767 | 0.743083 | 0.65093 | 0.39099 |
| | | | | 0.65128 |

DTA 120-200µm

| | CTL | | | | | |
|---|---|---|---|---|---|---|
| mouse1 | mouse2 | mouse3 | mouse4 | | mouse1 | mouse2 |
| 20 | 23.0769231 | 0 | 0 | | 25 | 26.6666667 |
| 33.3333333 | 25 | 6.25 | 15.3846154 | | 23.5294118 | 28.5714286 |
| 33.3333333 | 12.5 | 22.2222222 | 18.1818182 | | 54.5454545 | 25 |
| 4.54545455 | 33.3333333 | 33.3333333 | 28.5714286 | | 35.7142857 | 21.4285714 |

| Kir2.1 | |
| --- | --- |
| mouse3 | mouse4 |
| 7.69230769 | 16.6666667 |
| 18.1818182 | 30 |
| 12.5 | 33.3333333 |
| 44.4444444 | 16.6666667 |
